# Impaired AMPK Control of Alveolar Epithelial Cell Metabolism Promotes Pulmonary Fibrosis

**DOI:** 10.1101/2024.03.26.586649

**Authors:** Luis R. Rodríguez, Konstantinos-Dionysios Alysandratos, Jeremy Katzen, Aditi Murthy, Willy Roque Barboza, Yaniv Tomer, Rebeca Acín-Pérez, Anton Petcherski, Kasey Minakin, Paige Carson, Swati Iyer, Katrina Chavez, Charlotte H. Cooper, Apoorva Babu, Aaron I. Weiner, Andrew E. Vaughan, Zoltan Arany, Orian S. Shirihai, Darrell N. Kotton, Michael F. Beers

**Affiliations:** Pulmonary, Allergy, and Critical Care Division, Department of Medicine, Perelman School of Medicine at the University of Pennsylvania, Philadelphia, PA 19104, USA; PENN-CHOP Lung Biology Institute, Perelman School of Medicine at the University of Pennsylvania, Philadelphia, PA 19104, USA; Center for Regenerative Medicine, Boston University and Boston Medical Center, Boston, MA 02118, USA; The Pulmonary Center and Department of Medicine, Boston University Chobanian & Avedisian School of Medicine, Boston, MA 02118, USA; Departments of Medicine, Endocrinology and Molecular and Medical Pharmacology, David Geffen School of Medicine at UCLA, Los Angeles, CA 90095, USA; Department of Biomedical Sciences, School of Veterinary Medicine, University of Pennsylvania, Philadelphia, PA 19104, USA; Institute for Regenerative Medicine, University of Pennsylvania, Philadelphia, PA 19104, USA; Cardiovascular Institute, Perelman School of Medicine at University of Pennsylvania 19104 USA

**Keywords:** Idiopathic Pulmonary Fibrosis, Epithelial Reprogramming, Stem Cells, mitochondrial impairments, Alveolar Epithelial Cells, AMP Kinase

## Abstract

Alveolar epithelial type II (AT2) cell dysfunction is implicated in the pathogenesis of familial and sporadic idiopathic pulmonary fibrosis (IPF). We previously described that expression of an AT2 cell exclusive disease-associated protein isoform (SP-C^I73T^) in murine and patient-specific induced pluripotent stem cell (iPSC)-derived AT2 cells leads to a block in late macroautophagy and promotes time-dependent mitochondrial impairments; however, how a metabolically dysfunctional AT2 cell results in fibrosis remains elusive. Here using murine and human iPSC-derived AT2 cell models expressing SP-C^I73T^, we characterize the molecular mechanisms governing alterations in AT2 cell metabolism that lead to increased glycolysis, decreased mitochondrial biogenesis, disrupted fatty acid oxidation, accumulation of impaired mitochondria, and diminished AT2 cell progenitor capacity manifesting as reduced AT2 self-renewal and accumulation of transitional epithelial cells. We identify deficient AMP-kinase signaling as a key upstream signaling hub driving disease in these dysfunctional AT2 cells and augment this pathway to restore alveolar epithelial metabolic function, thus successfully alleviating lung fibrosis *in vivo*.

## INTRODUCTION

Idiopathic pulmonary fibrosis (IPF), an enigmatic chronic interstitial lung disease, claims the lives of more than 40,000 Americans every year^1^. The understanding of IPF shifted dramatically in the early 2000s moving from the pathophysiologic paradigm of an inflammation-driven disease, to that of aberrant epithelial wound healing resulting in progressive lung fibrosis^2–5^. Of the many types of epithelia present in the lung, a link between alveolar epithelial type II (AT2) cells in particular and progressive lung fibrosis was suggested by observations in histopathological studies of patients with IPF having prominent AT2 cell hyperplasia^6^ and enhanced AT2 cell apoptosis^7^. Within the alveolar niche, the AT2 cell represents a vital progenitor with the capacity to self-renew and differentiate into the AT1 cell to maintain alveolar epithelial integrity after injury^8–13^. Coupled with genetic studies demonstrating heritable interstitial lung disease-related variants in the AT2 cell-specific surfactant protein genes^14^, the concept of repetitive microinjuries in the alveolar space leading to AT2 cell dysfunction and initiation of the disease process emerged. More recently, several innovative preclinical models of pulmonary fibrosis have revealed pulmonary fibrosis arises as a result of dysfunctional AT2 cell endophenotypes characterized by markers of senescence^15^, ER stress^16,17^, disrupted macroautophagy^18–20^, telomere dysfunction^21,22^, and proinflammatory/profibrotic signaling^18,23^. Based on these studies, AT2 cell dysfunction has emerged as a driver of sustained fibrosis, rather than AT2 cell loss^11^ which results in a resolving fibrotic response^24,25^.

Understanding the pathobiology underlying IPF initiation and progression has been facilitated by the characterization of high effect size variants in the surfactant protein C (*SFTPC*) gene found in patients with familial pulmonary fibrosis (FPF), sporadic IPF, and childhood interstitial lung disease (chILD). Of particular relevance is the missense variant in the linker domain of the surfactant protein C (SP-C) pro-protein (*S*ftpc^I73T^) resulting in a mutant protein with altered intracellular trafficking patterns, including accumulation of misprocessed isoforms at the plasma membrane and within endosomal compartments^18,26^. *In vitro* studies using cell lines expressing SP-C^I73T^ revealed an acquired dysfunctional epithelial cellular phenotype marked by a late block in macroautophagy, impaired mitophagy, and defective cellular proteostasis^26^. In addition, AT2 cells from *S*ftpc^I73T^ mice demonstrate a similar phenotype accompanied by an early multiphasic injury/alveolitis (7-14 days) followed by a transition to fibrosis (14-42 days)^18^. In keeping with these observations, patient-specific *S*FTPC^I73T^ mutant iPSC-derived AT2 cells (iAT2s) display diminished AT2 cell progenitor capacity, proteostasis defects, mitochondrial impairments, and inflammatory activation^27^. Notably, these alterations in cellular metabolism and mitochondrial homeostasis are consistent across species and various *in vitro* and *in vivo* platforms. Thus, we hypothesized that a conserved epithelial metabolic phenotype underlies the pathobiology of fibrotic lung disease.

Disruption to homeostatic metabolic pathways, including oxidative phosphorylation and glycolysis, has been recognized as a component of the fibrotic milieu for more than a decade^28–31^. With several early studies focused on metabolic changes in fibrotic mesenchymal lineages^29,30,32–34^. Specifically, increased glycolysis with enhanced lactate generation was observed in activated fibroblasts and modulation of glycolytic enzymes inhibited fibrogenesis *in vitro* and in mice *in vivo*^33,35,36^. Alveolar epithelial mitochondrial impairments in IPF were first recognized in reports with the accumulation of enlarged, dysmorphic mitochondria with impaired ETC complex I and IV activity in AT2 cells from patients with IPF, a phenotype that was associated with decreased expression of the mitophagy regulator PINK1^37^. Not surprisingly, the defect in mitophagy was accompanied by impaired autophagy, with increased levels of autophagy markers LC3B and p62 in hyperplasic AT2 cells localized to fibrotic regions of IPF lungs^37^. Subsequently, alterations in mitochondrial homeostasis or activity have each been shown to impact AT2 cell function and lung fibrogenesis. For example, genetic disruption of mitochondrial networks in AT2 cells altered phospholipid and cholesterol metabolism^38^ and resulted in spontaneous lung fibrosis^38^. More recently, genetic manipulation of mitochondrial complex I activity was shown to alter AT2 cell progenitor function *in vivo*^39^ and defects in mitochondrial homeostasis of AT2 cells from IPF lungs and bleomycin injured mouse lungs were responsive to interventions involving thyroid hormone mimetics^40^.

Although these findings have greatly expanded our understanding of the role of AT2 cell-specific metabolic alterations in the pathobiology of IPF, the questions regarding how a metabolically dysfunctional AT2 cell leads to the initiation and progression of fibrosis and whether the restoration of alveolar epithelial function results in amelioration of established fibrosis remain elusive. Using murine and iPSC-derived AT2 cell models, we herein detail the altered mitochondrial dynamics, respiration, and proglycolytic state that emerge in AT2 cells as a result of SP-C^I73T^ expression and identify AMP-kinase (AMPK) as a druggable signaling hub to rescue the aberrant AT2 cell metabolic phenotype and downstream lung fibrosis.

## Results

### Murine AT2 Cells Expressing Sftpc^I73T^ Exhibit a Metabolic Shift Toward Glycolysis

The ontogeny of lung fibrosis in *S*ftpc^I73T^ mice following tamoxifen induction of mutant protein expression is marked by three distinct phases: initiation (days 0-3), inflammation (days 3-14), and fibrogenesis (days 14-28)^18,23^. To define the sequence of AT2 cell changes that occur during these epochs we performed population RNA sequencing (popRNA-seq) of AT2^I73T^ cells isolated at two key time points following *in vivo* tamoxifen administration: day 3 (“early” inflammation), day 14 (transition from inflammation to fibrogenesis), and AT2^WT^ cells. We identified 1921 and 2627 differentially expressed genes (DEGs) in day 3 and day 14 AT2^I73T^ cells, respectively, compared to AT2^WT^ cells (for a total of 3595 combined DEGs; **Supplemental Figure S1**). Based on STRINGDB^41^ and Enrichr^42^, genes associated with cell cycle regulation and extracellular matrix organization were differentially upregulated in AT2^I73T^ cells, whereas genes associated with multiple metabolic processes were differentially downregulated (**Figure 1A and Supplemental Figure S1**). Reactome pathway analysis identified differential regulation of multiple metabolic pathways including upregulation of carbohydrate and protein metabolism and downregulation of phospholipid and fatty acid metabolism pathways in AT2^I73T^ cells (**Figure 1B**).

**Figure 1:**
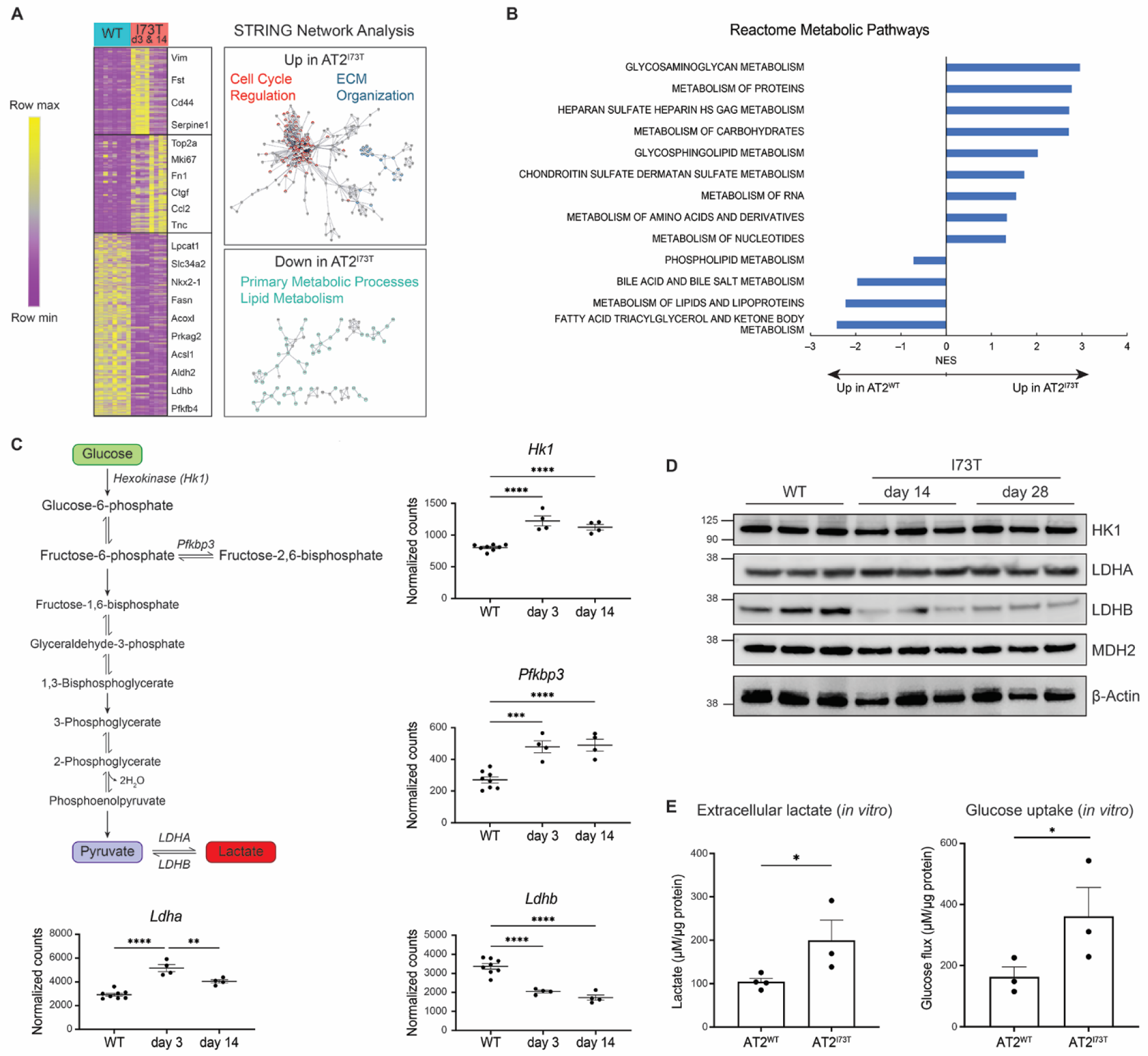
Increased Glycolysis in Murine AT2 Cells Expressing *S*ftpc*^I73T^*. A) Unsupervised hierarchical clustering using Euclidean distance heatmap of differential expression analysis (FC >1.5, FDR <0.05) comparing AT2^I73T^ cells 3- and 14-days post *in vivo* tamoxifen induction (n = 4 per group) to AT2^WT^ cells (derived from age matched C57/B6J mice, n = 12) by popRNA-seq. A subset of differentially expressed genes is highlighted. STRING network analysis demonstrating downregulation of genes associated with primary metabolic processes and lipid metabolism and upregulation of genes associated with proliferation and ECM organization in AT2^I73T^ cells. B) Reactome pathway analysis of differentially expressed genes in AT2^I73T^ cells 3- and 14-days post tamoxifen induction demonstrates differential regulation of multiple metabolic pathways. C) Graphical outline of rate-limiting enzymes in glycolysis pathway and individual graphs of normalized counts of day 3 and 14 AT2^I73T^, and AT2^WT^ cells by popRNA-seq for each highlighted gene. D) Western blot of AT2^I73T^ cells at peak of inflammation (day 14, n = 3) and fibrosis (day 28, n = 3) shows differential abundance of LDHA and LDHB proteins. E) Extracellular lactate and glucose concentrations (µM) in 48-hour *ex vivo* cultures of AT2^I73T^ cells isolated 28 days after in vivo tamoxifen administration and AT2^WT^ cells, measured by YSI biochemistry analyzer and normalized to total protein content. *p <0.05, **p <0.005, ***p <0.0005, ****p <0.00005 by ordinary one-way ANOVA.

In addition, examination of transcripts regulating glycolysis commitment steps in the glycolytic pathway showed upregulation of *Hk1* and *Pfkb3* in AT2^I73T^ cells while transcriptional shifts in the *Ldha/Ldhb* ratio also suggested increased generation of lactate from pyruvate (**Figure 1C**). Immunoblotting of AT2 cell lysates validated the differential abundance of LDHA and LDHB proteins **(Figure 1D**). Functionally, we observed increased glucose uptake and lactate production in short-term AT2^I73T^ cell cultures (**Figure 1E**). Collectively, these data suggest that murine AT2^I73T^ cells exhibit time-dependent metabolic alterations marked by an increased glycolytic state. Alterations to enzyme expression in the citric acid (TCA) cycle as well as differential levels of glycolytic and TCA cycle organic acids were also detected supporting additional functional effects on cell metabolism in AT2^I73T^(**Supplemental Figure S2**).

### Disrupted Mitochondrial Biogenesis and Altered Mitochondrial Functions in Murine AT2^I73T^ cells

The shift toward glycolysis that emerged following induction of *S*ftpc*^I73T^* was accompanied by significant alterations to the mitochondrial phenotype. Gene set enrichment analysis of Hallmark gene signatures^43^ based on the popRNA-seq analysis, identified a significant downregulation of the PPAR signaling pathway in AT2^I73T^ cells (**Figure 2A**), a pathway critical in the regulation of mitochondrial biogenesis. Furthermore, the majority of genes encoding mitochondria localized proteins (MitoCarta3.0 dataset^44^) were downregulated in AT2^I73T^ cells (**Figure 2A**). In support of this transcriptional signature, we observed a reduction in activated pPGC1α protein (**Figure 2B**) as well as a significant time-dependent reduction in mitochondrial DNA copy number (mtDNAcn) (**Figure 2C**), a marker of mitochondrial biogenesis^45,46^. Similarly, mtDNA gene expression was downregulated (**Figure 2D**) in AT2^I73T^ cells. In addition, we observed time-dependent reduction of mitochondrial membrane potential (ΔΨm) and respiration (basal, maximal, and spare respiratory capacity). These changes were observed as early as 14 days post *in vivo* induction of mutant SP-C expression (**Figure 2E, F**).

**Figure 2:**
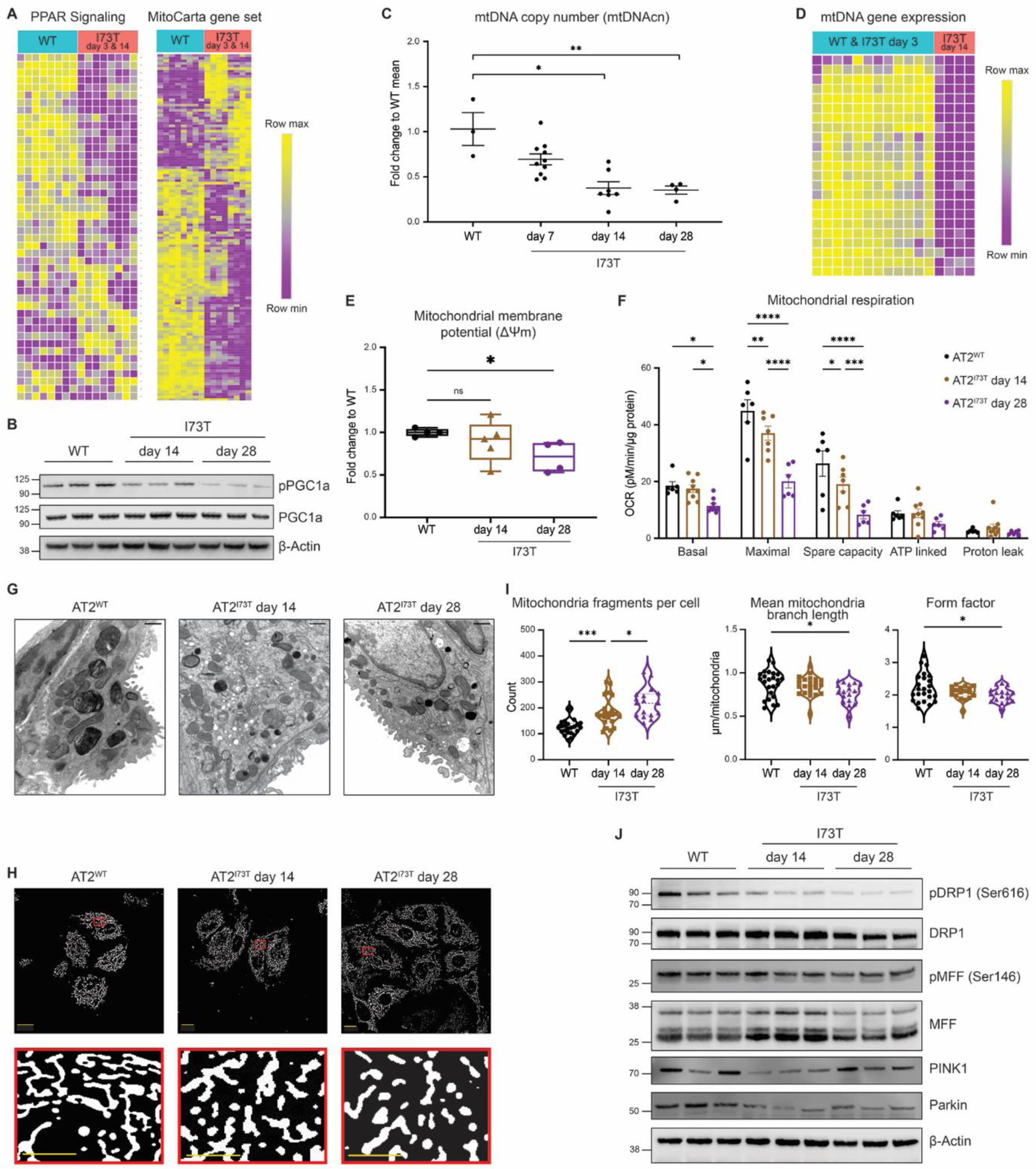
Defects in Mitochondrial Biogenesis and Mitochondrial Dynamics. A) Unsupervised hierarchical clustering using Euclidean distance heatmap of differentially expressed genes (FDR <0.05) in the Hallmark “PPAR Signaling” pathway and Mitocarta gene sets (row normalized z-score) demonstrates downregulation in AT2^I73T^ cells at 3- and 14-days post *in vivo* induction (n = 4 per time point). B) Western blot of activated phosphorylated PGC1α confirms decreased pathway activation (n = 3 per condition). C) Time-dependent reduction in mitochondrial DNA copy number (mtDNAcn) in AT2^I73T^ cells (7 and 14 days, n = 6; 28 days, n = 5). D) Unsupervised hierarchical clustering using Euclidean distance heatmap of differentially expressed mtDNA genes by popRNA-seq shows decreased expression in AT2^I73T^ cells 14-days post in vivo induction. E) Flow cytometry analysis of mitochondrial membrane potential (ΔΨm) in murine AT2 cells demonstrates a time-dependent reduction in Tetramethylrhodamine, methyl ester (TMRM) fluorescence in AT2^I73T^ cells (n = 4-5 per condition). F) Measurement of oxygen consumption rate (OCR) shows a time-dependent reduction in basal, maximal uncoupled mitochondrial respiration, and spare respiratory capacity in AT2^I73T^ cells (n = 4-9 per condition). G) Electron microscopy images of murine AT2 cells from whole lung mounts. Scale bars: 600 nm. H) Mitotracker staining of murine AT2 cells after 18 hours in culture shows altered mitochondrial network morphology in AT2^I73T^ compared to AT2^WT^ cells. Scale bars: 10nm. I) Quantification of mitochondrial structure in AT2 cells using ImageJ Mitochondrial Analyzer shows accumulation of mitochondrial fragments, decreased branch length, and altered shape in AT2^I73T^ cells. J) Western blot of key mitochondrial dynamics proteins demonstrates altered regulation of mitochondrial dynamics and mitophagy in AT2^I73T^ cells. *p <0.05, **p <0.005, ***p <0.0005, ****p <0.00005 by ordinary one-way ANOVA.

Given that functional impairments in mitochondria a (specifically ΔΨm alterations) are often matched by compensatory increases in mitochondrial clearance through mitophagy and that we have previously observed defects in autophagy and mitophagy *in vitro*^26,27^, we hypothesized that mitochondrial turnover could be compromised resulting in a reorganized AT2 cell mitochondrial network. Electron microscopy demonstrated significant ultrastructural alterations to the mitochondrial network of AT2^I73T^ cells (**Figure 2G**) manifested as an accumulation of smaller mitochondria, reminiscent of prior observations made in AT2 cells derived from IPF lungs^37^. Similarly, immunofluorescence microscopy of labeled mitochondria in isolated AT2 cells showed the presence of a disrupted mitochondrial network with more mitochondrial fragments per cell, decreased branch lengths, and altered fragment shape in AT2^I73T^ cells (**Figure 2H, I**). Next, we tested whether there was a link between the accumulation of fragmented dysfunctional mitochondria and disrupted mitochondrial turnover (**Figure 2J**). We observed decreased protein levels of the mitophagy regulators PINK1 and Parkin that in tandem with the loss of phosphorylated DRP1(Ser616) and MFF1(Ser146) suggested a DRP1-independent fission mechanism^47^. Of note, both DRP1 and MFF1 are targets of AMP-activated protein kinase (AMPK) with MFF1(Ser146) recognized as a canonical AMPK phosphorylation motif^48,49^. Taken together, these findings suggest diminished mitochondrial biogenesis and disruptions in ΔΨm, mitochondrial respiration, and mitochondrial fission/fusion dynamics in AT2^I73T^ cells.

### Impaired AMPK Signaling in murine AT2^I73T^ cells

To gain a mechanistic understanding of the relationship between disrupted AT2 cell metabolism, diminished mitochondrial biogenesis, and impaired macroautophagy, we focused on ATP levels and AMPK activity, a well characterized signaling hub regulating several cellular processes, including autophagy and mitochondrial homeostasis (**Figure 3A**)^50–53^. In line with the well-documented “Warburg effect”, initially observed in tumor cells and subsequently identified in some progenitor cell populations^53,54^, AT2^I73T^ cells displayed an early surge in ATP levels that were sustained through day 14 post-induction, followed by a return to baseline levels at the fibrotic 28-day time point (**Figure 3B)**. Next, we performed immunoblot analysis of AT2 cell lysates to interrogate AMPK signaling and its downstream regulation of fatty acid oxidation (FAO) through the acetyl-coA carboxylase (ACC) and observed decreased phosphorylation of AMPK with a concurrent decrease in ACC phosphorylation in AT2^I73T^ cells at 14 and 28 days post *in vivo* induction of mutant SP-C (**Figure 3C**).

**Figure 3:**
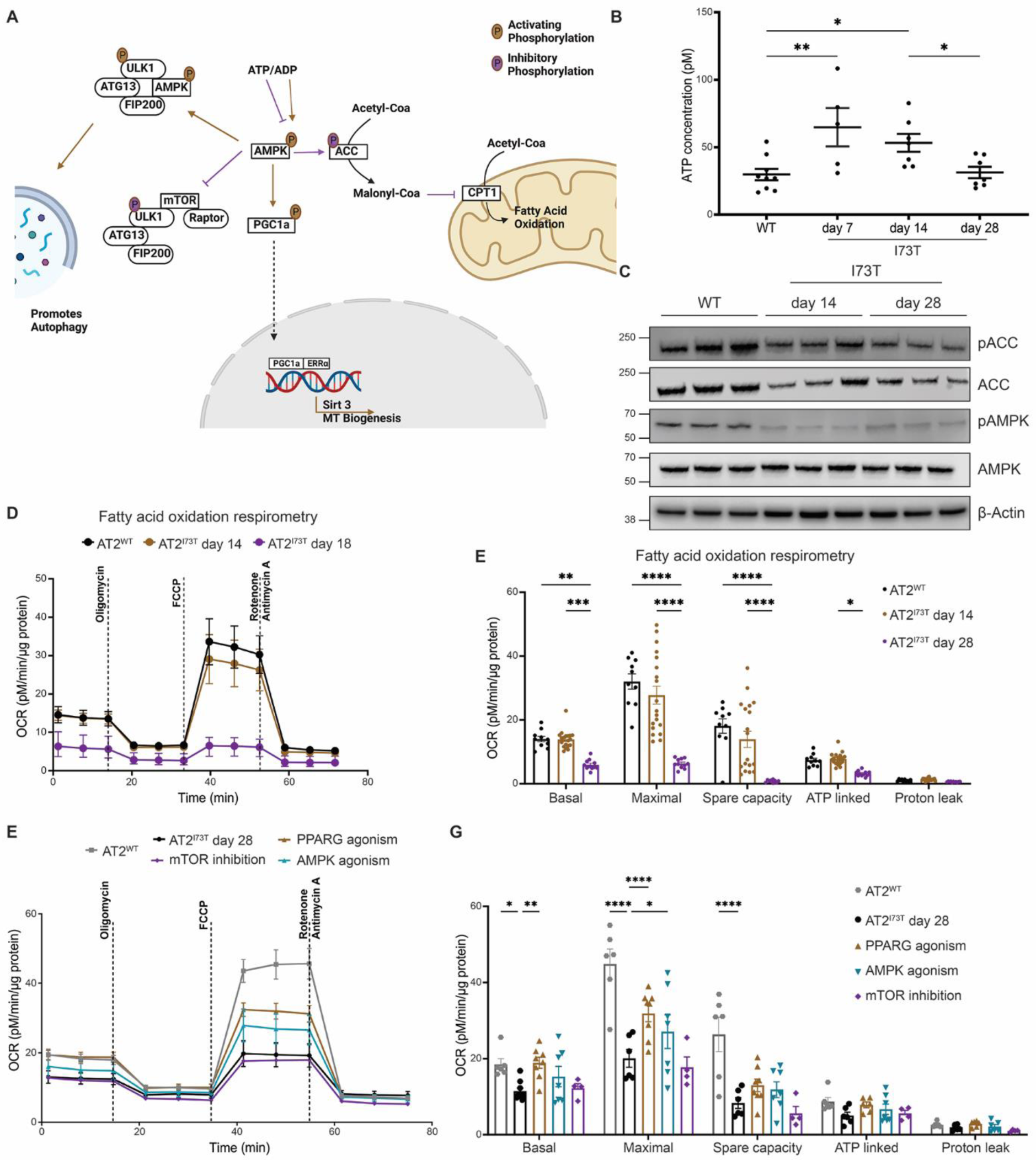
Impaired Fatty Acid Oxidation and Mitochondrial Respiration are linked to AMPK Signaling. A) Graphical representation of AMPK as a key regulator of many cellular processes including mitochondrial biogenesis, autophagy, and fatty acid oxidation (FAO). B) Increased accumulation of ATP in AT2^I73T^ cells freshly isolated 7- and 14-days post *in vivo* tamoxifen induction (25,000 cells, n = 5-7 per condition). C) Western blot analysis of key enzymes in AMPK signaling pathway demonstrates a decrease in AMPK signaling and FAO in AT2^I73T^ cells (n = 3 per condition). D,E) Reduced OCR in AT2^I73T^ cells isolated 14- and 28-days post *in vivo* tamoxifen induction and cultured overnight in media supplied with endogenous fatty acids (n = 9-15 per condition). F,G) OCR in AT2^I73T^ cells isolated 28 days post tamoxifen induction and cultured for 48 hours in the presence of the following small molecules: rosiglitazone (25 µM, n = 8) to stimulate PPARγ, PF-06409577 (100 nM, n = 7) to activate AMPK, and torin 1 (100 nM, n = 4) to inhibit mTOR. *p <0.05, **p <0.005, ***p <0.0005, ****p <0.00005 by ordinary one-way ANOVA.

To functionally assess the downstream consequences of disrupted AMPK signaling, particularly its influence on FAO, we analyzed the oxygen consumption of AT2 cells under minimal media conditions containing the long-chain fatty acid palmitate. We found significant time-dependent reductions in basal, maximal, and spare oxygen consumption rates (OCR) in AT2^I73T^ cells when limited to substrates promoting oxygen consumption through FAO (**Figure 3D, E**). These findings suggest that disrupted AMPK signaling not only stems from the altered metabolic phenotype but may also contribute to the dysfunction through regulation of downstream pathways including fatty acid metabolism.

We next tested potential approaches for reversing the bioenergetic deficiencies in AT2^I73T^ cells by modulating AMPK or its downstream pathways (**Figure 3F, G)**. We isolated AT2 cells from wild-type and *S*ftpc*^I73T^* mice 28 days post *in vivo* tamoxifen induction (fibrotic time point). AT2^I73T^ and AT2^WT^ cells were cultured in 2D and treated with: rosiglitazone to stimulate PPARγ, PF-06409577 to activate AMPK, and Torin 1 to inhibit mTOR. Cultured AT2^I73T^ cells maintained expression of mutant proSP-C and demonstrated reduced AMPK signaling as compared to AT2^WT^ cells (**Supplemental Figure 3**). Forty-eight hours of either direct PPARγ or AMPK activation in AT2^I73T^ cells rescued the mitochondrial respiration with significant increases in maximal OCR compared to vehicle controls, whereas mTOR inhibition had no effect on OCR (**Figure 3F, G**). Taken together, these findings suggest that the impaired AMPK signaling in AT2^I73T^ cells, associated with elevated ATP levels, results in defective FAO and mitochondrial respiration and that PPARγ activation or AMPK agonism effectively ameliorate the mitochondrial respiration defects.

### Diminished progenitor function and emergence of a transitional cell state resulting from Sftpc^I73T^ expression

In the intestinal epithelium, metabolic alterations have been associated with a defect in epithelial progenitor cell function^55^. To assess potential alterations in AT2^I73T^ progenitor cell function (self-renewal and differentiation to AT1 cells) due to the observed bioenergetic changes, we reanalyzed our recently published single cell RNA sequencing (scRNA-seq) time series profiles (GSE234604^56^) of *S*ftpc^I73T^ mouse lungs (14 and 28 days post tamoxifen induction; **Supplemental Figure 4**). Within this dataset composed of 35002 cells, 2500 alveolar epithelial cells were identified and reclustered for further analysis (**Figure 4A and Supplemental Figure 4)**. Louvain clustering identified 5 alveolar cell clusters. Based on top DEGs and published canonical markers, two of these were annotated as AT2 cells^57–59^ (cluster 2 and 4), one as AT1 cells (cluster 1), and two as putative transitional cells (clusters 3 and 5; detailed below) (**Figure 4B and Supplemental Figure 5)**. KEGG pathway analysis of the DEGs defining each cluster identified PPAR signaling as the most upregulated pathway in cluster 2 AT2 cells (hereafter designated AT2-PPAR High; **Figure 4C, 4D and Supplemental Figure 5**), whereas PPAR signaling was lower in cluster 4 AT2 cells (hereafter AT2-PPAR Low). AT2-PPAR High cells were also distinguished from AT2-PPAR Low cells by significantly lower AMPK signaling, biosynthesis of unsaturated fatty acids, and mitophagy (**Supplemental Figure 5H, 5I).**

**Figure 4:**
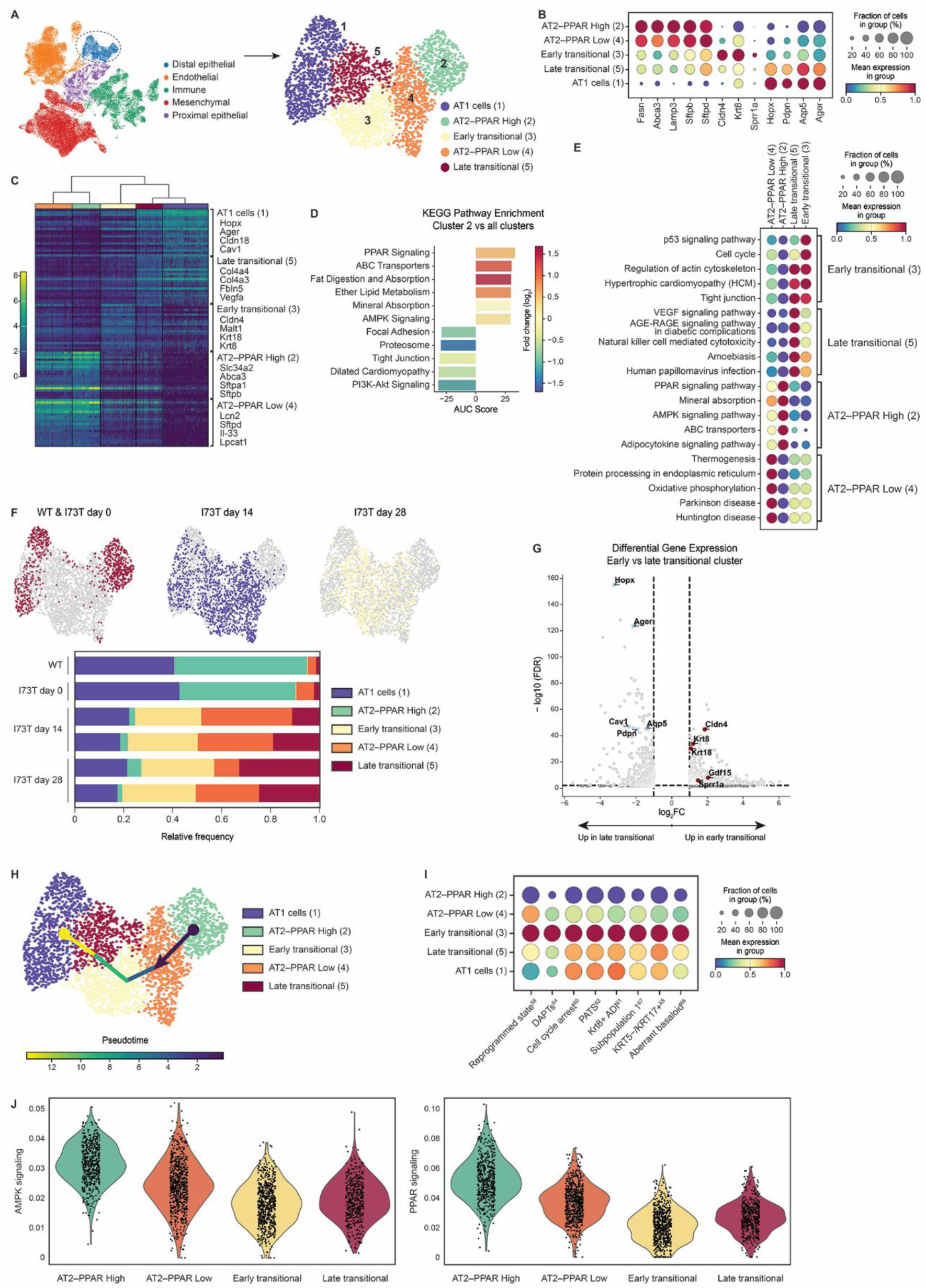
Metabolic alterations and emergence of an epithelial transitional state in response to *in vivo S*ftpc*^I73T^* expression. A) Uniform manifold approximation and projection (UMAP) visualization of 35002 lung cells profiled by scRNA-seq in GSE234604 color-coded by cell lineage with subset analysis of 2500 distal epithelial cells. B) Gradient dot plot of key distal epithelial genes used to annotate AT1 (1), AT2 (2, 4), and transitional subclusters (3, 5). C) Dendrogram of top 20 DEGs and their associated log2FC for each cluster as determined by false discovery rate (FDR). A subset of cluster-defining genes is highlighted. D) KEGG Pathway enrichment analysis of DEGs (FDR <0.05, log2FC>1 and <-1) in cluster 2 as compared to all other clusters. E) Gradient dot plot of KEGG Pathway enrichment analysis performed on differentially upregulated genes (FDR <0.05, log2FC>1) for each cluster as compared to all other clusters. F) Color-coded UMAPs by genotype and time point and frequency table denoting the frequency of each cluster within biological samples. G) Volcano plot of differential expression analysis (FDR <0.05, log2FC>1 or <- 1) comparing the early transitional cluster 3 to the late transitional cluster 5 highlighting decreased expression of AT1 cell marker genes (*Hopx*, *Ager*, *Cav1*, *Pdpn*, *Aqp5*) and increased expression of transitional cell marker genes (*Cldn4*, *Krt8*, *Krt18*, *Gdf15*, *Sppr1a*) in the early transitional cell cluster. H) Pseudotime trajectory analysis with starting node set in the AT2-PPAR High cluster. I) Gradient dot plot demonstrating expression of indicated transitional state gene modules (Supplemental Table 1) across the distal epithelial clusters in the day 28 *S*ftpc^I73T^ time point. J) KEGG Pathway module scores of AMPK and PPAR signaling demonstrate progressively reduced expression from the AT2-PPAR High cluster to the transitional cell clusters.

In lung injury models, disrupted AT2-to-AT1 differentiation was recently linked to the emergence of a putative “transitional state”, characterized by a gene signature that in mice includes *Krt8* and *Cldn4*^58,60,61^. Given that clusters 3 and 5 were enriched in transitional state markers, including *Krt8, Krt18, Sox4, Fn1*, and *Cldn4* (**Figure 4B**), we annotated these as transitional cell clusters. We noted the relative abundance of cluster 3 remained unchanged between the day 14 and day 28 time points, whereas cluster 5 was more abundant in the day 28 time point (**Figure 4F**). Comparing cluster 3 to cluster 5, we observed the differential upregulation of classical AT1 cell marker genes (*Hopx*, *Pdpn*, *Ager*, *Cav1*, *Aqp5*) in cluster 5 (**Figure 4G**). Furthermore, pseudotime analysis using a starting node in the AT2-PPAR High cluster suggested that cluster 3 serves as an early precursor state to cluster 5 (**Figure 4H and Supplement Figure 5**). Based on these observations, we chose to refer to cluster 3 as “early” and cluster 5 as “late” transitional cluster.

To further contextualize the 2 putative transitional states, we next defined transitional cell gene modules based on recently published datasets of diseased mouse or human lungs that included scRNA-seq profiles of these cells, variably referred to as PATS^62^, cell cycle arrest^63^, DATPs^64^, KRT5−/KRT17+^65^, aberrant basaloid^66^, Krt8+ alveolar differentiation intermediate (ADI)^61^, subpopulation 1^67^, or reprogrammed state^58^ (**Supplemental Table 1**). Of all our annotated cell clusters, we found early transitional cells (cluster 3) had the highest expression levels of all transitional cell modules defined from prior reports, irrespective of the injury model or species (**Figure 4I**). Furthermore, using AMPK signaling and PPAR signaling pathway gene modules we observed a progressively decreased expression of these pathways as AT2^I73T^ cells enter the transitional state (**Figure 4J**).

Next, we used immunofluorescence (IF) microscopy on paraffin embedded sections from *S*ftpc^I73T^ mouse lungs at 14- and 28-days post tamoxifen induction of mutant SP-C^I73T^, as well as WT mouse lungs, to localize KRT8^high^ cells. Consistent with a recent study in a model of bleomycin lung injury^68^, we found the emergence of KRT8^high^ proSP-C co-expressing cells <u>prior</u> to widespread fibrogenesis (day 14) and the persistence of that cell state through the fibrotic stage (day 28), where KRT8^high^ proSP-C+ cells localized to areas of ASMA^+^ injury (**Figure 5A**). Using a novel flow cytometry approach (**Supplemental Figure 6**), we confirmed the IF staining findings by measuring an increased frequency of transitional cells as fibrosis developed (**Figure 5B**). Furthermore, flow cytometry analysis confirmed the loss of AT1 cells (**Figure 5C**) suggested by the scRNA-seq quantification (**Figure 4F**). Single cell suspensions of AT2, transitional, and AT1 cells isolated from *S*ftpc^I73T^ mouse lungs 14 days post *in vivo* tamoxifen induction, revealed a gradual reduction in MitoTracker^TM^ fluorescence from AT2 to transitional to AT1 cells, suggesting a gradual loss of mitochondria mass during the AT2-to-AT1 differentiation (**Figure 5D),** consistent with AT2 gene expression profiles (**Figure 2**).

**Figure 5:**
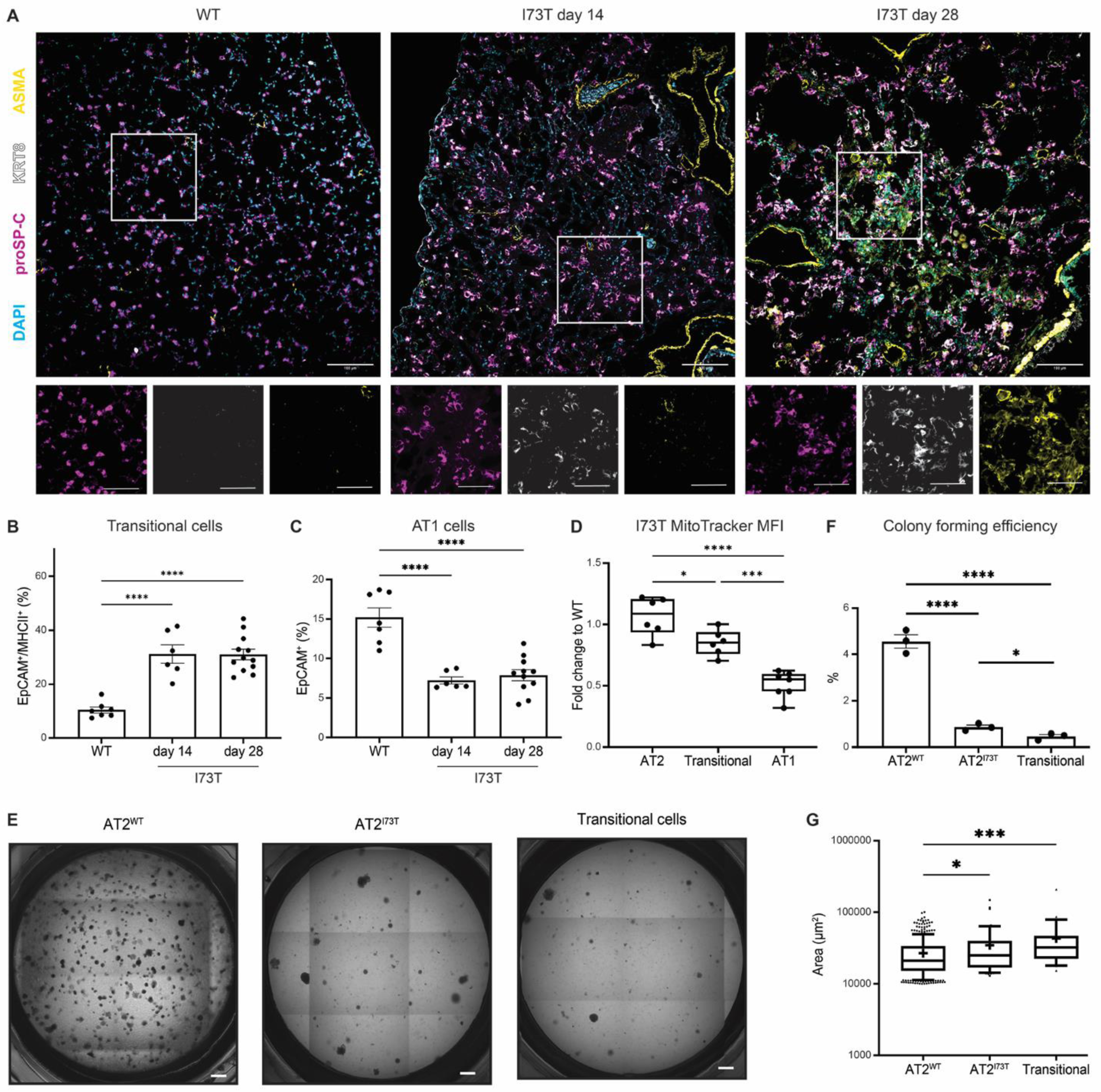
Persistence of an epithelial transitional state with diminished progenitor capacity in *S*ftpc*^I73T^*lungs. A) Representative immunofluorescent staining of murine lung sections derived from WT or *S*ftpc*^I73T^* mice 14- and 28-days post tamoxifen induction stained with antibodies against proSP-C, KRT8, ASMA, and DAPI for nuclei counterstaining. Scale bars: 100 µm. B) Flow cytometry quantification of CD51+ transitional cells demonstrate a sustained increase of transitional cells after *in vivo* tamoxifen induction (n = 6-12 per time point). C) Flow cytometry quantification of EpCAM+/CD51/-/CD104-AT1 cells demonstrates a sustained loss of AT1 cells after *in vivo* tamoxifen induction (n = 6-12 per time point). D) Mean fluorescence intensity (MFI) of MitoTracker® dye accumulating in AT2, transitional, and AT1 cells isolated from *S*ftpc*^I73T^* mice 14 days post *in vivo* tamoxifen induction (n = 6-7 per time point). E) Representative light microscopy images of 21-day organoid cultures derived using WT PDGFRα^+^ fibroblasts and 1) AT2 cells isolated from WT mice, 2) AT2^I73T^ cells, and 3) transitional cells isolated from *S*ftpc*^I73T^* mice 14 days post *in vivo* tamoxifen induction. Scale bars: 500 µm. F) Colony forming efficiency (CFE) of organoids with surface area >>10,000 µm^2^ (n = 3 per condition). G) Surface area quantification of organoids with surface area > 10,000 µm^2^. *p <0.05, ***p <0.0005, ****p <0.00005 by ordinary one-way ANOVA.

We assessed the capacity of AT2^I73T^ cells to self-renew, a measure of their progenitor potential. We established three-dimensional (3D) organoid cultures using WT PDGFRα^+^ fibroblasts and 1) AT2 cells isolated from WT mice, 2) AT2^I73T^ cells, and 3) transitional cells (**Supplemental Figure 6**) isolated from *S*ftpc*^I73T^*mice 14 days post *in vivo* tamoxifen induction. After 14 days in culture, colony forming efficiency (CFE) was assessed (**Figure 5E**). We observed a significant decrease in CFE for AT2^I73T^ and transitional cells compared to AT2^WT^ cells, with transitional cells exhibiting the lowest CFE (**Figure 5F**). This was consistent with our previous findings in human patient-specific SFTPC^I73T^ expressing iAT2s^27^. Although AT2^I73T^ and transitional cells formed fewer organoids, these were larger in size as compared to the AT2^WT^ cell-derived organoids (**Figure 5G**). Collectively these data support diminished AT2 cell progenitor capacity as a result of mutant SP-C^I73T^ expression, evident as reduced AT2 self-renewal potential coincident with reduced presence of differentiated AT1 cells and accumulation of transitional cells.

### AMPK Agonism Ameliorates the Metabolic Alterations Observed in human iPSC-derived AT2^I73T^ cells

To test whether human AT2 cells expressing SP-C^I73T^ exhibit similar perturbations, we next employed a second preclinical *in vitro* platform of human AT2 cell dysfunction that enables testing of epithelial intrinsic effects without the secondary effects arising from other cell lineages. We recently reported the generation of iPSCs from patients with *S*FTPC*^I73T^* variants and lung disease^27^ and identified that expression of mutant SP-C^I73T^ in iAT2s results in diminished AT2 cell progenitor function, autophagy perturbations, altered bioenergetic programs, time-dependent increased glycolysis, and activated canonical NF-kB signaling^27^. Immunoblot analysis of iAT2^I73T^ cells vs syngeneic cells gene-edited to express normal (WT) SFTPC iAT2^WT^ cells revealed a quantitative increase in the LDHA/LDHB ratio in iAT2^I73T^ cells (**Figure 6A, B**). Similar to mouse AT2^I73T^ cells (**Figure 1)** and consistent with our recently reported increase in extracellular acidification rate (ECAR)^27^, human iAT2^I73T^ cells demonstrated increased lactate production (**Figure 6C)**. Reanalysis of our previously published popRNA-seq dataset^27^ for expression of transcripts associated with FAO, glycolysis, and oxidative phosphorylation further confirmed metabolic alterations in iAT2^I73T^ cells (**Supplemental Figure 7**), paralleling the changes identified in murine AT2^I73T^ cells (**Figure 1A-D**).

**Figure 6:**
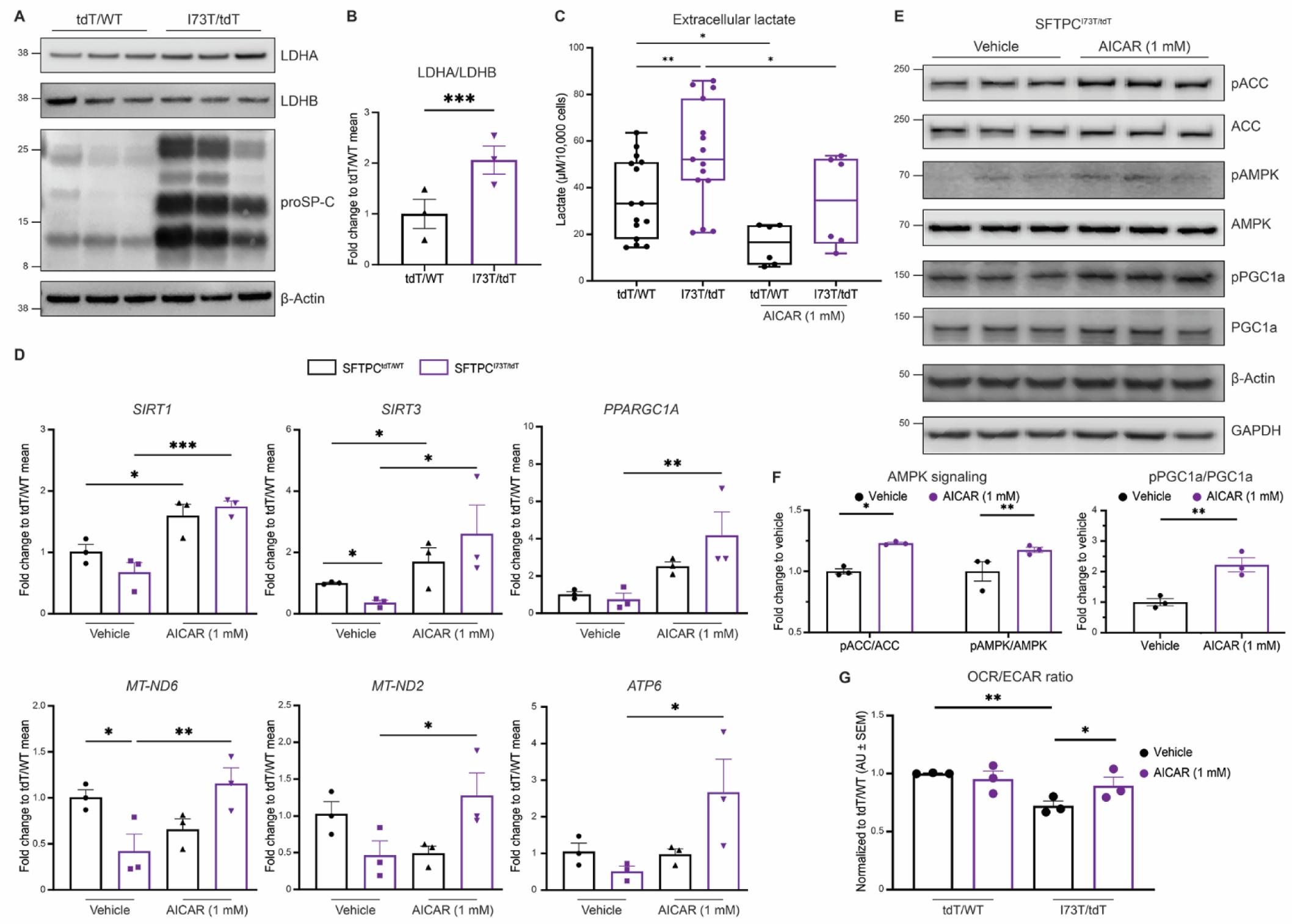
AMPK Agonism Ameliorates the Metabolic Alterations Observed in human iPSC-derived AT2^I73T^ (iAT2s^I73T^) cells. A-B) Western blot analysis of serially cultured human iAT2 cells (135-182 days) demonstrates increased LDHA/B ratio in iAT2^I73T^ (*S*FTPC*^I73T/tdTomato^*) cells compared to syngeneic corrected iAT2s (*SFTPC^WT/tdTomato^*). C) Measurement of extracellular lactate by YSI biochemistry analyzer shows increased extracellular lactate in iAT2^I73T^ cells compared to corrected iAT2s. This increase is significantly reduced by treatment with AICAR (1 mM) for 24 hours (n = 9 per condition). D) RT-qPCR of PGC1α target genes demonstrates increased mitochondrial biogenesis in iAT2^I73T^ cells after treatment with AICAR (1 mM) for 24 hours (n = 3 per condition). E, F) Western blot of AMPK pathway targets in iAT2^I73T^ cell lysates after treatment with AICAR (1 mM) for 24 hours confirms activation of AMPK signaling and increased phosphorylation of PGC1α. Bar graphs depict densitometric quantification (n = 3 per condition). G) Respirometry quantification depicted as OCR/ECAR ratio in iAT2^I73T^ and corrected iAT2 cells following treatment with AICAR (1 mM) for 24 hours. *p <0.05, **p <0.005, ***p <0.0005, by ordinary one-way ANOVA.

Given that AMPK activation in mouse AT2^I73T^ cells in vitro alleviated the mitochondrial respiration defects, we sought to determine whether the increased glycolysis and reduced mitochondrial biogenesis (**Figure 6D**) observed in human iAT2^I73T^ cells could be ameliorated through direct AMPK stimulation. We treated iAT2^I73T^ cells with AICAR 1mM for 24 hours, a dose and duration confirmed to stimulate AMPK signaling (**Supplemental Figure 7**). AICAR treatment led to a reduction in extracellular lactate levels in both mutant and corrected iAT2s, consistent with a reduction in glycolytic shift (**Figure 6C**) and corrected the acquired defects in autophagy (**Supplemental Figure 7B-D**) and NF-kB signaling (**Supplemental Figure 7E)**. Gene expression analysis of transcripts associated with mitochondrial biogenesis, coupled with immunoblot analysis, confirmed that AMPK agonism via AICAR increased the phosphorylation of PGC1α and enhanced mitochondrial biogenesis, including increased expression of multiple PGC1α target genes (**Figure 6D-F**). Functionally, the improved mitochondrial biogenesis observed in human iAT2^I73T^ cells following AICAR treatment was accompanied by enhanced mitochondrial respiration (**Figure 6G**). Collectively, these findings suggest that direct AMPK agonism in human iAT2^I73T^ cells through AICAR enhances mitochondrial biogenesis and ameliorates the observed mitochondrial respiration defects.

### AMPK Agonism Promotes AT2 Cell Respiration in vivo and Rescues the Fibrotic Sftpc^I73T^ Lung Phenotype

Given that AMPK agonism reversed the AT2^I73T^ cell metabolic phenotype in two distinct *in vitro* platforms, we then sought to test whether a similar strategy might effectively reverse the observed metabolic changes *in vivo* as a therapeutic approach. In a clinically relevant interventional dosing strategy (**Figure 7A**), metformin administration (150 mg/kg given intraperitoneally 5 out of 7 days a week) initiated on day 12 post tamoxifen induction of mutant SP-C^I73T^ expression resulted in significantly improved survival from less than 70% in the vehicle treated mice to 90% in the intervention arm (p =0.0199, **Figure 7B**). Gross histological examination of lung structure in surviving mice at 28 days revealed a qualitative improvement characterized by a decreased cellular infiltrate and improved alveolar architecture with administration of metformin (**Figure 7C**). Quantitatively, these changes were reflected in improvements in lung physiology, measured as static lung compliance (**Figure 7D, E**), and reductions in total cell counts in bronchoalveolar lavage fluid (BALF) and total BALF protein (**Figure 7F, G**). The effect size of these metrics compares favorably with previously published data demonstrating the efficacy of nintedanib (BIBF1120) in the *S*ftpc^I73T^ mouse model^56^.

**Figure 7:**
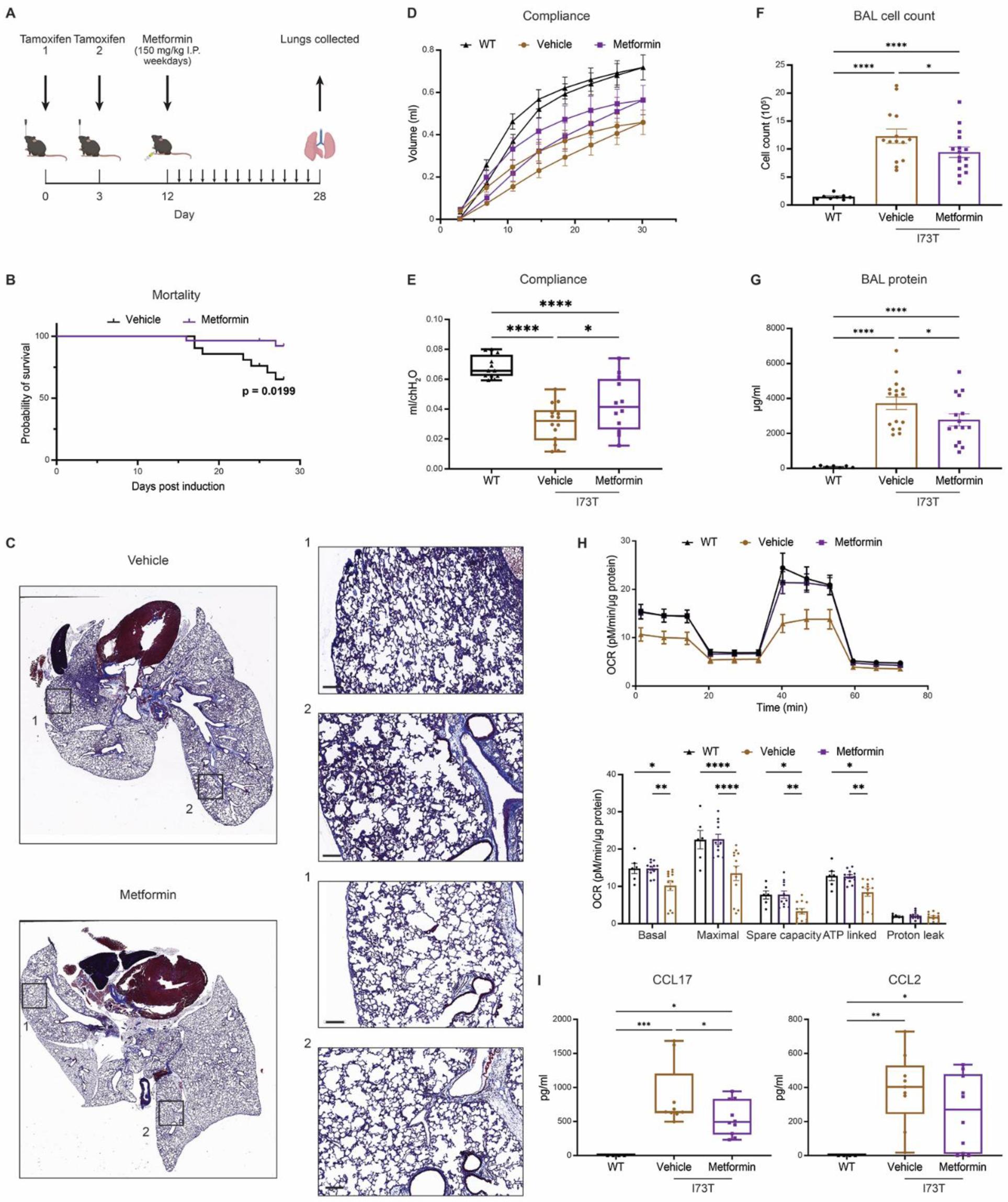
*In Vivo* Rescue of the Sftpc^I73T^ Fibrotic phenotype Via Metformin Intervention. A) Study design for metformin intervention in the *S*ftpc*^I73T^*mouse fibrosis model. Delivery of intraperitoneal (i.p.) metformin (150 mg/kg) or vehicle control was performed daily on weekdays, beginning at day 12 post tamoxifen induction (n = 20 per group). B) 28-day mortality was significantly (p=0.0199) reduced in metformin-treated mice. C) Histology of lungs collected from mice at 28 days post tamoxifen induction and treated with either metformin or vehicle control. D, E) Measurement of static lung compliance via Scirec Flexivent demonstrates a significant increase with metformin treatment as compared to vehicle control (n = 12 per group). F, G) Markers of inflammation and lung injury in the bronchoalveolar lavage (BAL) fluid collected from mice at 28 days post tamoxifen are significantly reduced in mice treated with metformin compared to the vehicle control (n = 12 per group). H) OCR measurement and quantification in AT2 cells isolated from WT and *S*ftpc*^I73T^* mice 28 days post *in vivo* tamoxifen induction. AT2 cells were seeded overnight before respirometry was performed (n = 8 per group). I) ELISA quantification of CCL17 and CCL2 concentrations in BAL fluid of WT and *Sftpc^I73T^* mice 28 days post *in vivo* tamoxifen induction. *p <0.05, **p <0.005, ***p <0.0005, ****p <0.00005 by ordinary one-way ANOVA.

To test whether the observed improvement in the fibrotic phenotype is linked to the correction of the dysfunctional AT2 cell metabolic signature, we isolated AT2^I73T^ cells from *S*ftpc*^I73T^* mice treated with either vehicle or metformin 28 days post *in vivo* tamoxifen induction and evaluated OCR. Similar to the *in vitro* findings (**Figure 3F, G**), *in vivo* AMPK agonism resulted in improved AT2 cell mitochondrial respiration evident as improvements in basal, maximal, ATP-linked, and spare respiratory capacity (**Figure 7H**). Metformin treatment also reduced BALF levels of CCL17 and CCL2, two cytokines we previously reported as being upregulated in AT2^I73T^ cells and increased in the *S*ftpc*^I73T^* mouse model of lung fibrosis^18^ (**Figure 7I**). Taken together, these findings suggest that *in vivo* AMPK agonism, achieved through metformin administration, ameliorates lung fibrosis in the *Sftpc^I73T^* mouse model and enhances AT2 cell mitochondrial function.

## DISCUSSION

Through *in vitro* and *in vivo* models we identified four key metabolic alterations in PF associated mutant AT2^I73T^ cells: increased glycolysis, impaired mitochondrial biogenesis, disrupted mitochondrial respiration, and decreased fatty acid oxidation. We further identified AMPK as a critical metabolic signaling hub deficient in AT2^I73T^ cells and found that AMPK activation can resolve the observed AT2 metabolic alterations as well as the broader fibrotic lung phenotype in *Stfpc^I^*^73T^ mice. Collectively, our results highlight a previously unrecognized role of epithelial metabolism in distal lung homeostasis.

Initial studies describing metabolic dysfunction in IPF were primarily focused on the mesenchymal compartment and the role of lactate in fibroblast activation^29,69^. Inhibiting LDHA was shown to be effective *in vitro* and in preclinical models *in vivo* in limiting fibrotic endpoints^69,70^. In our study, we found elevated lactate production in murine AT2^I73T^ and human iAT2^I73T^ cells, with changes in LDH subunit composition, consistent with prior *ex vivo* observations made using AT2 cells obtained from end-stage human IPF lungs^71^. The generation of lactate observed in our models under aerobic conditions (**Figure 1E**), coupled with altered transcription of glycolytic enzymes (**Figure 1C**), increased glucose flux (**Figure 1E**), changes in FAO (**Figures 2-3**), impaired oxidative phosphorylation (OXPHOS) (**Figure 2F**), and accumulation of cellular ATP content (**Figure 4B**) aligns with the emerging concept of a non-cancerous Warburg effect as an important regulator of cell state^54,72^. Effective maintenance of the alveolar epithelium, whether during normal cell homeostasis or in response to injury, relies on effective AT2 cell progenitor functions of self-renewal and differentiation to AT1 cells. We found that AT2^I73T^ cells had impaired mitochondrial respiration (**Figures 2F and 6G**) and increased glycolysis (**Figures 1C-E and 6A-C**) consistent with a pathological metabolic state with diminished progenitor function, both *in vivo* **(Figures 4F and 5A-C)** and *in vitro* (**Figure 5E,F and Alysandratos et al.** ^27^). An association between disrupted epithelial cell respiration and pulmonary fibrosis has been previously reported in Pink1^-/-^ mice exhibiting impaired bioenergetics and age-related spontaneous fibrosis^37,40^. In two mouse models of pulmonary fibrosis, thyroid hormone administration improved mitochondrial biogenesis and bioenergetics leading to amelioration of pulmonary fibrosis in an extrinsic murine injury model (bleomycin), an effect mediated through PGC1α and PINK1^40^. Similarly, genetic disruption of mitochondrial networks has been associated with impaired epithelial energy production and the promotion of spontaneous fibrosis^38^. However, the precise signals by which disruptions in cellular bioenergetics and metabolism impact AT2 cell function, ultimately leading to pulmonary fibrosis, remained elusive.

Our observations provide a mechanistic link between an aberrant AT2 cell metabolic phenotype and fibrogenesis suggested by the impaired AMPK signaling in AT2^I73T^ cells (**Figure 3**) and the diminished AT2^I73T^ cell progenitor capacity, evident as reduced AT2 cell self-renewal in organoid models coincident with reduced presence of AT1 cells and accumulation of epithelial transitional cells in areas of fibrosis in our mouse model (**Figures 4 and 5**). The identification of a transitional cell state in multiple lung injury models^58,61–67^ has raised new questions regarding the role and relevance of this putative cell state to disease pathogenesis and our findings suggest that metabolic dysfunction may play a role in the emergence of this state.

Our scRNA-seq analysis further reinforced the role of mitochondrial biogenesis and AMPK signaling (**Figure 4J**) in this fibrotic model and prompted us to consider the therapeutic implications of AMPK agonism. Previous studies focusing on fibroblast biology recognized AMPK as a druggable target for addressing metabolic dysfunction in IPF^32,73–75^. Furthermore, there is now ample evidence indicating that stimulating mitochondrial biogenesis and enhancing mitochondrial respiration in AT2 cells can improve outcomes in mouse models of lung fibrosis^34,40,76,77^. To explore this therapeutic potential in IPF, we harnessed the iAT2^I73T^ cell platform for both target identification and validation^27^. Leveraging the reductionist nature of this model, we first confirmed the altered glycolytic phenotype and consequent increase in lactate production (**Figure 6A-C**). Given that the inherent defect in the model lies in the autonomous generation of mutant SP-C^I73T^, we can deduce that the initial metabolic phenotype is a direct response to mutant protein expression, rather than a secondary response to “outside-in” signaling. Utilizing this model, we also identified that AMPK agonism restored mitochondrial biogenesis and stimulated mitochondrial respiration (**Figure 6D-G**). The use of the iAT2 cell platform confirms and underscores the multifaceted role of AMPK as a key hub for AT2 cell homeostasis. Furthermore, these findings provide supporting evidence that a comprehensive approach targeting multiple metabolic pathways can be a valid strategy, laying the foundation for the *in vivo* application of AMPK agonism.

To provide proof of concept for the *in vitro* findings, we chose metformin, an indirect activator of AMPK with a reliable pharmacokinetic profile in mice, FDA approval for the treatment of metabolic disorders^78^, and emerging epidemiological evidence for clinical benefits in IPF^79^. Consistent with prior data generated in a bleomycin model of pulmonary fibrosis^32^, we observed significant improvements in survival, lung physiology, and markers of injury and immune infiltration (**Figure 7B-G**). While we cannot exclude the broad targeting of mesenchymal and immune populations associated with disease progression, the rescue of AT2 cell respiration and reduction in cytokines produced by AT2 cells during fibrosis, suggest that metformin’s beneficial effects are, at least in part, mediated through the amelioration of AT2 cell metabolic dysfunction. Taken together, our findings affirm the efficacy of AMPK agonism in rescuing AT2 cell metabolism and mitigating aberrant fibrogenesis, further linking these events to the resolution of AT2 cell dysfunction in the *Sftpc^I73T^*mouse *in vivo*.

In conclusion, our data support a role for epithelial metabolic dysfunction in IPF mediated by an AMPK hub and contribute to the growing body of evidence that metabolic intervention stands as a viable approach for the treatment of pulmonary fibrosis.

## AUTHOR CONTRIBUTIONS

DNK and MFB developed the concept. LRR, KDA, JK, DNK, and MFB designed the experiments. YT, PC, SI, KC, CHC performed *in vivo* animal experiments. LRR, KDA, AM, RAP, AP, KM, PC, SI, KC, CHC conducted experiments and analyzed data. WRB and AB performed bioinformatic analysis. AIW, AEV, JK, AM, and LRR conceived, validated, and optimized flow cytometry strategies. LRR, KDA, JK, RAP, AP, OSS, DNK, and MFB interpreted data and generated figures. LRR and KDA drafted the original manuscript. LRR, KDA, JK, WRB, RAP, AP, AIW, AEV, ZA, OSS, DNK, and MFB edited the manuscript. All authors reviewed and approved the final version prior to submission.

## ACKNOWLEDGMENTS

The authors wish to thank all members of the Beers, Kotton, and Alysandratos Labs for insightful discussions. Michael F. Beers is an Albert M. Rose Established Investigator of the Pulmonary Fibrosis Foundation and is the Robert L. Mayock and David A. Cooper Professor of Medicine. We thank the PENN-CHOP Lung Biology Institute Informatics team for guidance. Multiple figures were created using Biorender.com.

## METHODS

### Sftpc^I73T^ Mouse Model of Pulmonary Fibrosis

Tamoxifen inducible *S*ftpc^I73T/I73T^ Rosa26ERT2FlpO^+/+^ (a.k.a. I^ER^-*S*ftpc*^I73T^*) mice expressing an NH2-terminal HA-tagged murine *S*ftpc*^I73T^* mutant allele into the endogenous mouse *Sftpc* locus were previously generated as reported^18^ and are summarily detailed. Tamoxifen treatment of adult I^ER^-*S*ftpc*^I73T^* mice was initiated at 12-14 weeks of age by oral gavage (OG) on day 0 and day 3. All mouse strains and genotypes generated for these studies were congenic with C57/B6/J. Both male and female animals (aged 8-14 weeks) were utilized in tamoxifen induction protocols. All mice were housed under pathogen free conditions in an AALAC approved barrier facility at the Perelman School of Medicine, University of Pennsylvania. All experiments were approved by the Institutional Animal Care and Use Committee at the University of Pennsylvania.

### iPSC line generation and maintenance

The SPC2 iPSC line clones SPC2-ST-C11 and SPC2-ST-B2 were used in this study. As previously detailed^27^, we used TALENs to insert a tdTomato fluorescent reporter at the translation initiation (ATG) site of the endogenous *SFTPC* locus of the parental SPC2 iPSC line, resulting in the generation of either corrected (SPC2-ST-B2 clone; SFTPC^tdT/WT^) or mutant (SPC2-ST-C11 clone; SFTPC^I73T/tdT^) iPSC clones as the tdTomato cassette is followed by a stop/polyA cassette, preventing expression of the subsequent *SFTPC* coding sequence from the targeted allele. iPSCs used in this study demonstrated a normal karyotype when analyzed by G-banding and/or array Comparative Genomic Hybridization (aCGH, Cell Line Genetics). iPSCs were maintained in feeder-free conditions, on growth factor-reduced Matrigel (Corning) in 6-well tissue culture dishes (Corning), in mTeSR1 media (StemCell Technologies) using gentle cell dissociation reagent for passaging. Further details of iPSC derivation, characterization, and culture are available for free download at https://crem.bu.edu/cores-protocols/#protocols.

### iPSC-directed Differentiation into Alveolar Epithelial Type 2 Cells (iAT2s) and Maintenance

To generate iAT2s, we performed PSC-directed differentiation via definitive endoderm into NKX2-1 lung progenitors using methods we have previously described^27,80–82^. On day 15 or 16 of differentiation, live cells were sorted on a high-speed cell sorter (MoFlo Astrios EQ) to isolate NKX2-1+ lung progenitors based on CD47^hi^CD26^−^ gating^82^. Sorted lung progenitors were resuspended in undiluted growth factor-reduced 3D Matrigel (Corning) at a density of 400 cells/μl, and distal/alveolar differentiation of cells was performed in CK+DCI medium, consisting of complete serum-free differentiation medium base supplemented with 3 μM CHIR99021, 10 ng/mL recombinant human KGF (CK), and 50 nM dexamethasone (Sigma), 0.1 mM 8-Bromoadenosine 3′,5′-cyclic monophosphate sodium salt (Sigma), and 0.1 mM IBMX (Sigma) (DCI). The resulting epithelial spheres were passaged without further sorting on approximate day 30 of differentiation followed by a brief period (4-5 days) of CHIR99021 withdrawal to achieve iAT2 maturation, as previously shown^80^. After this 2-week period, SFTPC^tdTomato+^ cells were purified by fluorescence activated cell sorting (FACS) to establish pure cultures of iAT2s. iAT2s were then maintained through serial passaging as self-renewing monolayered epithelial spheres (“alveolospheres”) by plating in 3D Matrigel (Corning) droplets at a density of 400 cells/μl with refeeding every other day with CK+DCI medium, according to our published protocol^81^. iAT2 culture quality and purity were monitored at each passage by flow cytometry, with > 90% of cells expressing SFTPC^tdTomato^ over time, as we have previously detailed^80,81^.

### Multichannel Flow Cytometry for Identification of Lung Cell Populations

Flow cytometry was performed as we have previously described^18,23,58^. Blood free perfused lungs were digested in Phosphate Buffered Saline (Mg and Ca free) with Collagenase Type I (Gibco Cat# 17100017), DNase (Millipore Sigma Cat# D5025), and Dispase (BD Biosciences). Resulting product was passed through 70-μm nylon mesh to obtain single-cell suspensions, and then processed with ACK Lysis Buffer (Thermo Fisher). Cells were incubated with antibody mixtures (see **Key Resources Table**). Stained cells were analyzed with an LSR Fortessa (BD Biociences). AT2 cells were identified as EpCAM+, CD45-, CD31-, MHCII^hi^, CD104-. Transitional AT2 cells were defined as EpCAM+, CD45-, CD31-, MHCII^hi^, CD104-, CD51^hi^. Quantification of mitochondrial mass was performed using MitoTracker™ dye and mitochondrial membrane potential was quantified using MitoProbe™ TMRM Assay Kit for Flow Cytometry according to manufacturer instructions. Cell populations were defined, gated, and analyzed with FlowJo software (FlowJo, LLC, Ashland, Oregon).

### Isolation of Mouse AT2 Cells

Mouse AT2 cells were isolated as previously reported^18,58,83^. Briefly, perfused mouse lungs were digested as described for flow cytometry to obtain a single cell suspension. Negative selection of mesenchymal cells via differential adherence on plastic culture dishes was followed by CD45 depletion using simultaneous incubation with Dynabeads untouched mouse T cell kit (Thermo Fisher #11413D) and Dynabeads mouse DC enrichment kit (#11429D) (Thermo Fisher Scientific). Subaliquots were analyzed for purity (greater than 90%) by flow cytometry using EpCAM+, proSP-C+ staining.

### Lung Histology

Whole lungs were fixed by tracheal instillation of 10% neutral buffered formalin (MilliporeSigma) at a constant pressure of 25 cm H2O. 6 μM sections were stained with Hematoxylin & Eosin (H&E) or Masson’s Trichrome stains by the Pathology Core Laboratory of Children’s Hospital of Philadelphia. Slides were scanned using an Aperio ScanScope Model: CS2 (Leica) at 10X magnification and representative areas captured and exported as TIF files and processed in Adobe Illustrator.

### Bronchoalveolar Lavage Fluid (BALF) Collection, Processing, and Cytokine Measurement

BALF collected from mice using sequential lavages of lungs with five X 1 ml aliquots of sterile saline was processed for analysis as previously described^18^. First ml of saline wash is labeled as enriched cell free BALF used to measure total protein and cytokines. Cell pellets obtained by centrifuging BALF samples at 400 g for 6 minutes were re-suspended in 1 ml of PBS, and total cell counts determined using a NucleoCounter (New Brunswick Scientific, Edison, NJ). Differential cell counts were determined manually from BALF cytospins stained with modified Giemsa (Sigma Aldrich, ^#^GS500) to identify macrophages, lymphocytes, eosinophils and neutrophils. Total protein content of cell free BALF was determined using the DC Protein Assay Kit (Cat ^#^ 5000111; BioRAD, Inc, Hercules CA) with bovine serum albumin as a standard according to the manufacturer’s instructions.

Total CCL2 and CCL17 concentration in the enriched cell free BALF was calculated using mouse CCL2 and CCL17 DuoSet Elisa (R&D Systems) according to the manufacturer’s instructions.

### Measurement of Pulmonary Function

At takedown, mice underwent Flexivent (SCIREQ, Inc. Toronto Canada) analysis for assessment of lung physiology as previously described^17,84^. Briefly, invasive measurement of static lung compliance was performed with mice anesthetized with intraperitoneal pentobarbital. The mouse tracheas were cannulated with a 20-gauge metal stub adapter and then placed on a small-animal ventilator at 150 breaths per min and a tidal volume of 10 ml/kg of body weight. Static lung compliance was determined with the manufacturer’s software using a 2 second breath pause maneuver.

### Mitochondrial immunofluorescence, confocal microscopy, and quantitation

AT2s were isolated as described above and seeded on cover slips via an adapted previously published protocol^85^. Briefly, cover slips were coated with 20% Matrigel (Corning) and 80% rat tail collagen (Gibco) for 2 hours at 37 prior to seeding with freshly isolated murine AT2s in 10% DMEM overnight. Cells were then stained using Mitotracker™ red according to manufacturer instructions. After staining, cells were washed and incubated for an additional 4 hours in 10% DMEM to allow for recovery of mitochondrial networks. Cells were then fixed using ice cold methanol followed by counterstaining using DAPI. Coverslips were imaged under confocal microscopy at 60X magnification using a Leica Stellaris 5 confocal. Images were quantified in Image J using the mitochondrial analyzer suite^86^.

### Electron Microscopy

Preparation of lung tissue and acquisition of transmission electron microscopy (TEM) images of lung sections was performed in the Electron Microscopy Resource Laboratory in the Perelman School of Medicine based on the method of Hayat that includes post-fixation in 2.0% osmium tetroxide with 1.5% potassium ferricyanide, as previously published^18^. Cut thin sections (60-80 nm) were stained in situ on copper grids with uranyl acetate and lead citrate and examined with a JEOL 1010 electron microscope fitted with a Hamamatsu digital camera and AMT Advantage image capture software.

### RNA Isolation and Quantitative Real Time Polymerase Chain Reaction (RT-qPCR)

RNA was extracted by first lysing cells in Qiazol (Qiagen) and subsequently using RNAeasy mini kit (Qiagen) according to the manufacturer’s protocol. The concentration and quality of extracted RNA from the lung tissues were measured using NanoDrop® One (Thermo Scientific, Wilmington, DE) and reverse-transcribed into cDNA using Verso cDNA Synthesis Kit (ThermoFisher). RT-qPCR was performed on a QuantStudio 7 Flex Real-Time PCR System with results normalized to *18S* and *Actb* gene expression. Primer sequences for all mouse and human genes are listed in **Key Resources Table**.

### Cellular ATP Quantification

AT2 cells were isolated as described above. ATP was measured using the Firefly Luciferase ATP Assay (Millipore Sigma) according to manufacturer instructions. Briefly, immediately after isolation 100,000 AT2 cells were mixed with d-luciferin, luciferin, and lysis solution in a 96-well plate. Luminescence was then measured using a TECAN Spark® multimode plate reader.

### In-vitro Culture of AT2 cells and Measurement of Metabolites

AT2 cells were isolated as described above and seeded in 10% DMEM for up to 48 hours. Validation of mutant surfactant protein C expression was performed via immunoblot analysis. Supernatants were collected after 24 hours in culture followed by measurement of glucose and lactate concentrations using YSI 2500 biochemical analyzer and according to manufacturer instructions.

### Immunoblot Analysis

iAT2s were harvested by incubating with 2 mg/ml dispase (Thermo Fisher Scientific) for 30-60 minutes at 37°C, treated with lysis buffer (RIPA buffer, 1x Roche Complete Protease Inhibitor cocktail, 1X Sigma Phosphatase Inhibitor Cocktail 2, and 1X Sigma Phosphatase Inhibitor Cocktail 3), and incubated on ice for 30 minutes. AT2 cells were either collected immediately after isolation or 48 hours after culture and treated with lysis buffer. Cellular debris was cleared by centrifugation at 15,000 g for 20 minutes and supernatants were harvested. Protein concentration was measured using Bio-Rad DC Protein Assay. 5-30 µg of lysates were resolved on pre-cast 10% or 12% Bis-Tris NuPAGE gels (Invitrogen), transferred to PVDF membranes (Bio-Rad), and blotted with primary antibodies overnight followed by 1 hour species specific secondary antibody incubation. All antibodies are listed in **Key Resources Table**. Visualization was performed on the Odyssey Imaging System (LiCOR Biosciences).

### Population RNA Sequencing

Mouse AT2 cells were collected, and RNA extracted as above. Library prep was performed by GeneWiz, LLC. Fastq files were evaluated for quality control with the FastQC program and then aligned against the mouse reference genome (mm10) using the STAR aligner^87^. Duplicate reads were flagged with the MarkDuplicates program from Picard tools and excluded from analysis. Per gene read counts for Ensembl (v67) gene annotations were computed using the R package Rsubread. Gene counts, represented as counts per million (CPM), were nominalized using TMM method in the edgeR R package, and genes with 25% of samples with a CPM < 1 were considered low expressed and removed. The data were transformed with the VOOM function from the limma R package to generate a linear model and perform differential gene expression analysis^88^. We employed the empirical Bayes procedure as implemented in limma to adjust the linear fit and to calculate P values given the small sample size of the experiment. We adjusted P values for multiple comparisons using the Benjamini-Hochberg procedure. Heatmaps were generated using Morpheus (https://software.broadinstitute.org/morpheus). Protein-protein interactions were obtained using STRING (Search Tool for Retrieval of Interacting Genes/Proteins) database. Gene ontogeny analysis was performed using the Database for Annotation, Visualization, and Integrated Discovery (https://david.ncifcrf.gov/home.jsp) based on differentially enriched genes (FC >1.5 P-value <0.05). Key pathway analyses was performed on gene lists identified from the GSEA molecular signature database^43,89,90^.

A previously published popRNA-seq dataset^27^ comparing corrected (SFTPC^tdT/WT^) and mutant (SFTPC^I73T/tdT^) iAT2s at day 131 of differentiation was re-analyzed in this manuscript. Experimental triplicate (n=3) samples of purified RNA extracts were harvested from each group of samples. Sequencing of pooled libraries was done using a NextSeq 500 (Illumina) and sequence reads were aligned to a combination of the human genome reference (GRCh38) and the tdTomato reporter sequence, using STAR v.2.5.2b^87^. Counts per gene were summarized using the featureCounts function from the subread package v.1.6.2. The edgeR package v.3.25.10 was used to import, organize, filter and normalize the data. Genes that were not expressed in at least one of the experimental groups were filtered out keeping only genes that had at least 10 reads in at least 3 libraries. The TMM method was used for normalization. The limma package v3.39.19^88,91^ with its voom method, namely, linear modelling and empirical Bayes moderation was used to test differential expression (moderate t tests). P values were adjusted for multiple testing using Benjamini-Hochberg correction (FDR). Differentially expressed genes between the groups in each experiment were visualized using Glimma v 1.11.1, and FDR <0.05 was set as the threshold for determining significant differential gene expression. Differential expression was then mined for genes that were annotated by the Camera package with KEGG and Reactome pathway enrichment terms containing metabolism or biosynthesis. Genes annotated with these terms were then process through Morpheus and STRING as above.

### Analysis of Single Cell RNA Sequencing

Distal epithelial cell specific analysis of our previously published data set (GSE234604) is included in this manuscript. Single cell RNA-Seq reads were aligned to mouse genome (mm10/GRCm38) using STARSolo (version 2.7.5b). After initial quality control and processing, we analyzed the scRNA-seq data using the Scanpy pipeline^92^. Genes expressed in fewer than 3 cells were removed, and cells with fewer than 200 genes and a mitochondrial fraction of less than 20% were excluded. Counts were log-normalized using scanpy.pp.normalize_per_cell (counts_per_cell_after=1x10^4^), followed by scanpy.pp.log1p. To integrate data from multiple samples, we used Scvi-tool^93^. We applied scvi.model.SCVI.setup_anndata() to establish the model parameters for integration, including: layer, categorical_covariate_keys, and continuous_covariate_keys. We then performed a principal component analysis (PCA) and generated a K-nearest neighbor (KNN) graph using scanpy.pp.neighbors with n_neighbors=15. The resulting KNN graph was used to perform Uniform Manifold Approximation and Projection (UMAP) dimension reduction to visualize the cells in two dimensions using scanpy.tl.umap(). Clustering was performed using the Leiden algorithm with scanpy.tl.leiden^94^. We identified cell populations using known canonical marker genes or by assessing cluster-defining genes based on differential expressions. Epithelial cells were clustered into proximal and distal clusters as reported in Supplemental Figure 7. Additionally, we performed linear trajectory inference on the UMAP reduction using scFates^95^ with the AT2 cluster as the starting point and without assigned endpoints. Finally, we performed gene ontology analysis for enriched biological processes using GSEApy^96^ based on differentially enriched genes between the groups.

### Cellular Respirometry assays

Human iPSC-derived alveolospheres were harvested by incubating with 2 mg/ml dispase (Thermo Fisher Scientific) for 30-60 minutes at 37°C, washed, and resuspended in 150-300 µl of Seahorse XF Base Media Minimal DMEM (Agilent Technologies) containing 2.8 mM glucose and 0.1 % FBS (pH 7.4). Alveolospheres were seeded in an XF96e Seahorse plate as described^27,97,98^. Briefly, growth factor reduced Matrigel (1.5 µl/well; Corning) was first pipetted in the center measurement zone of each well. The alveolosphere suspension (5-10 µl) was deposited in the Matrigel-coated measurement zone using a pipette. The plate was then incubated in a non-CO2 incubator for 3.5 minutes to let the Matrigel solidify. Next, 150 µl of pre-warmed Seahorse media was slowly added to each well to avoid dislodging the organoids from the Matrigel. The plate containing the organoids was centrifuged at 500 g for 5 minutes with no brake and subsequently incubated in a non-CO2 incubator for 30-45 minutes prior to running the assay. Oxygen consumption was determined as described^97,98^. Port injections were as follows: port A, oligomycin (final concentration 4.5 μM/l); port B, FCCP diluted in a mixture of 80% sodium pyruvate and 20% of 1:1 l-leucine/l-glutamine (final concentration 1 μM/l FCCP in 11.4 mM/l sodium pyruvate and 2.9 mM/l each of leucine/glutamine); and port C, antimycin A (final concentration 2.5 μM/l). Once the Seahorse assay was completed, the plate was kept for mitochondrial content determination. Protein concentration was determined by BCA assay (Thermo Fisher Scientific).

Primary mouse AT2s were isolated and plated as described above with measurement of mitochondrial respiration performed using the Seahorse XF Cell Mito Stress Test Kit (Agilent) or the Seahorse XF Palmitate Oxidation Stress Test Kit (Agilent) according to the manufacturer’s instructions. Prior to the initiation of the assay cells were maintained in either DMEM or, for small molecule challenge, under the various small molecules described in the results and figure legends. As with alveolospheres, stress test conditions included 2.5 µg/ml oligomycin, 2 µM FCCP, and 0.5 µM rotenone/antimycin A. Protein concentration was determined using DC protein assay (Bio-Rad).

### Primary Mouse Organoid Culture

Organoid culture assays were performed as previously described^58^. AT2 cells were flow sorted as described above, GFP positive fibroblasts were collected via flow cytometry from B6.129S4-Pdgfratm11(EGFP)Sor/J mice. In each technical replicate, 5000 AT2 cells were combined with 50,000 *Pdgfra*^+^ lung fibroblasts in 50% Matrigel (Corning) and 50% SAGM (Lonza) in a Falcon Cell Culture Insert. Cell/matrigel suspension solidified and SAGM medium was then added into the bottom of the well. SAGM was prepared according the BulletKit (Lonza) manufacturer instructions with some modification (Hydrocortisone, BSA, Triiodothyronine, and Epinephrine were not included). Medium was changed every other day. 10 μM rock inhibitor (Y-27632 dihydrochloride, Millpore Sigma, catalog # Y0503) was added to the medium for the first two days of culture. Organoids were imaged on an EVOS FL Microscope and were quantified via ImageJ using analyze particles macro.

### Statistics

All data are presented with dot-plots and group mean ± SEM unless otherwise indicated. Statistical analyses were performed with GraphPad Prism (San Diego, CA). Two tailed Student’s t-test were used for 2 groups as indictated; Multiple comparisons were performed by analysis of variance (ANOVA) with post hoc testing as indicated; survival analyses was performed using Kaplan Meier with Mantel Cox correction. In all cases statistical significance was considered at p values ≤ 0.05.

### Data and code availability

The sequencing data generated in this study are deposited in Gene Expression Omnibus (GEO) and will be made available upon publication of this work. Analysis of previously published data was performed on GSE234604.

### Study Approval

Mice housed in pathogen free facilities were subjected to experimental protocols approved by the IACUC of the Perelman School of Medicine at the University of Pennsylvania.

