## Supplemental Data for "Impaired AMPK Control of Alveolar Epithelial Cell Metabolism Promotes Pulmonary Fibrosis"

**Running Title:** *Epithelial AMPK Signaling in Lung Fibrosis*

### Lead Contact Author:

Michael F. Beers[†], M.D.
Pulmonary and Critical Care Division
Perelman School of Medicine at The University of Pennsylvania
Edward J Stemmler Hall Suite 216
3450 Hamilton Walk
Philadelphia, Pennsylvania 19104-6118


[†] Albert M. Rose Established Investigator of the Pulmonary Fibrosis Foundation

**Conflict of Interest:** The authors declare that no conflicts of interest exist.

| Supplemental Table 1: Transition Cell Gene Modules | | | | | | |
|---|---|---|---|---|---|---|
| PATS | Cell Cycle Arrest | DATPs | Krt5-Krt17+ | Aberrant Basaloid | Krt8+ | Subpopulation 1 |
| S100a6 | Cdk4 | Cldn4 | Krt17 | Cdh1 | Sprr1a | Cldn4 |
| Sfn | Trp53 | Krt8 | Prss2 | Cdh2 | Cldn4 | Sprr1a |
| Tmsb10 | Cdk2b | Cdkn1a | Krt7 | Spink1 | Cdkn1a | Tnip3 |
| Cldn4 | Ccnd1 | Ndrg1 | Gdf15 | Mmp7 | Plaur | Plaur |
| Clu | Aqp5 | Sprr1a | Mmp7 | Ptgs2 | Tnip3 | Prss23 |
| AW112010 | Emp2 | Tnip3 | Sox4 | Cdkn2a | Tnfrsf12a | AW112010 |
| Krt19 | Nfkbia | Mif | Tacstd2 | Cdkn2b | Edn1 | Cyr61 |
| Krt18 | Fn1 | Pold4 | Sfn | Hmga2 | Tmp2 | Fn1 |
| Anxa1 | Gadd45b | Mboat1 | Ociad2 | Epcam | S100a6 | Tpm2 |
| Krt8 | Serpine1 | Hif1a | Mdk | Vim | S100a10 | Bok |
| Tpm2 | Ctgf | Pdk4 | S100a2 | Fn1 | Anxa1 | Camp |
| Krt7 | Pdgfb | Cxx1b | Itgb6 | Col1a1 | Prkcdbp | Clu |
| Serpinb9 | Tgif1 | Lrrc26 | Tm4sf1 | Cdh2 | S100a14 | Itgb6 |
| Anxa2 | Itgb6 | Cdkn2a | Krt8 | Tnc | Lgals3 | Ngp |
| Ifitm3 | Nfib | Mdm2 | Krt18 | Vcan | Cyr6l | F3 |
| Epcam | Itgav | Ccnd1 | Tpm1 | Pcp4 | Atf3 | Prkcdbp |
| Lgals3 | Tgfb2 | Gdf15 | Ceacam6 | Cux2 | Krt18 | Sfn |
| Tnip3 | Tgfbi | Trp53 | Krt19 | Prss2 | Cryab | Anxa1 |
| Sox4 | | Bax | Pcsk1n | Cpa6 | Sfn | Ly6a |
| Anxa5 | | Infgr1 | Ptgs2 | Ctse | Rap2b | Fxyd3 |
| Cyr61 | | Ly6a | Ctse | Mdk | Myl12a | Tubb2b |
| Cavin3 | | Irf7 | Hopx | Gdf15 | Rps27l | Areg |
| Tuba1a | | Cxcl16 | Col1a1 | Slco2a1 | Tpm1 | Krt19 |
| S100a11 | | Timp1 | C8orf4 | Ephb2 | Anxa3 | Tnfrsf12a |
| Serpinb1a | | | Gprc5a | Itgb8 | S100a11 | Gsto1 |
| Crip1 | | | Pon2 | Itgav | Krt8 | S100a14 |
| Lgals1 | | | Lamb3 | Itgb6 | Anxa5 | Krt7 |
| Ccl20 | | | Cldn4 | Tgfbi | Anxa2 | Tuba1a |
| Actn1 | | | Tram1 | Kcnn4 | Cd81 | Malt1 |
| Mfge8 | | | Epcam | Kcnq5 | Cstb | Ceacam1 |
| Ubd | | | Fhl2 | Kcns3 | Krt7 | Emp3 |
| Tubb5 | | | Itga2 | Cdkn1a | Ubb | Eno1 |
| Ywhah | | | Ccnd2 | Ccnd1 | Epcam | Krt18 |
| S100a10 | | | Tagln | Ccnd2 | Hspb1 | Krt8 |
| Hspb1 | | | Phlda2 | Mdm2 | Ccng1 | Hbegf |
| 2200002D01Rik | | | Cst6 | Hmga2 | Msn | S100a6 |
| Nfkbia | | | Ccnd1 | Ociad2 | Rpl19 | Mfge8 |
| Cyba | | | Tnc | Ptchd4 | Rps5 | Cks2 |
| F3 | | | Sdc1 | | Rplp0 | S100a8 |
| Cd24a | | | Cdkn2a | | Sqstm1 | Cxcl2 |
| Cd81 | | | C19orf33 | | Igfbp7 | St3gal4 |
| Lurap1l | | | Lamc2 | | Calm2 | Scgb3a2 |
| Myl12a | | | Pcp4 | | Eef1a1 | Atf3 |
| S100a14 | | | Lbh | | Rps19 | Ddx39 |
| Tpm1 | | | Pmepa1 | | Pkm | Thbs1 |
| Igfbp7 | | | Zfp36l1 | | B2m | Ier5 |
| Marcksl1 | | | Cdh1 | | Arpc2 | Ly6d |
| Cfl1 | | | Icam1 | | Esd | Slc26a4 |
| | | | Tnfrsf12a | | Hsp90ab1 | |
| | | | Cd24 | | Cystm1 | |
| | | | | | Clu | |
| | | | | | Psmd8 | |

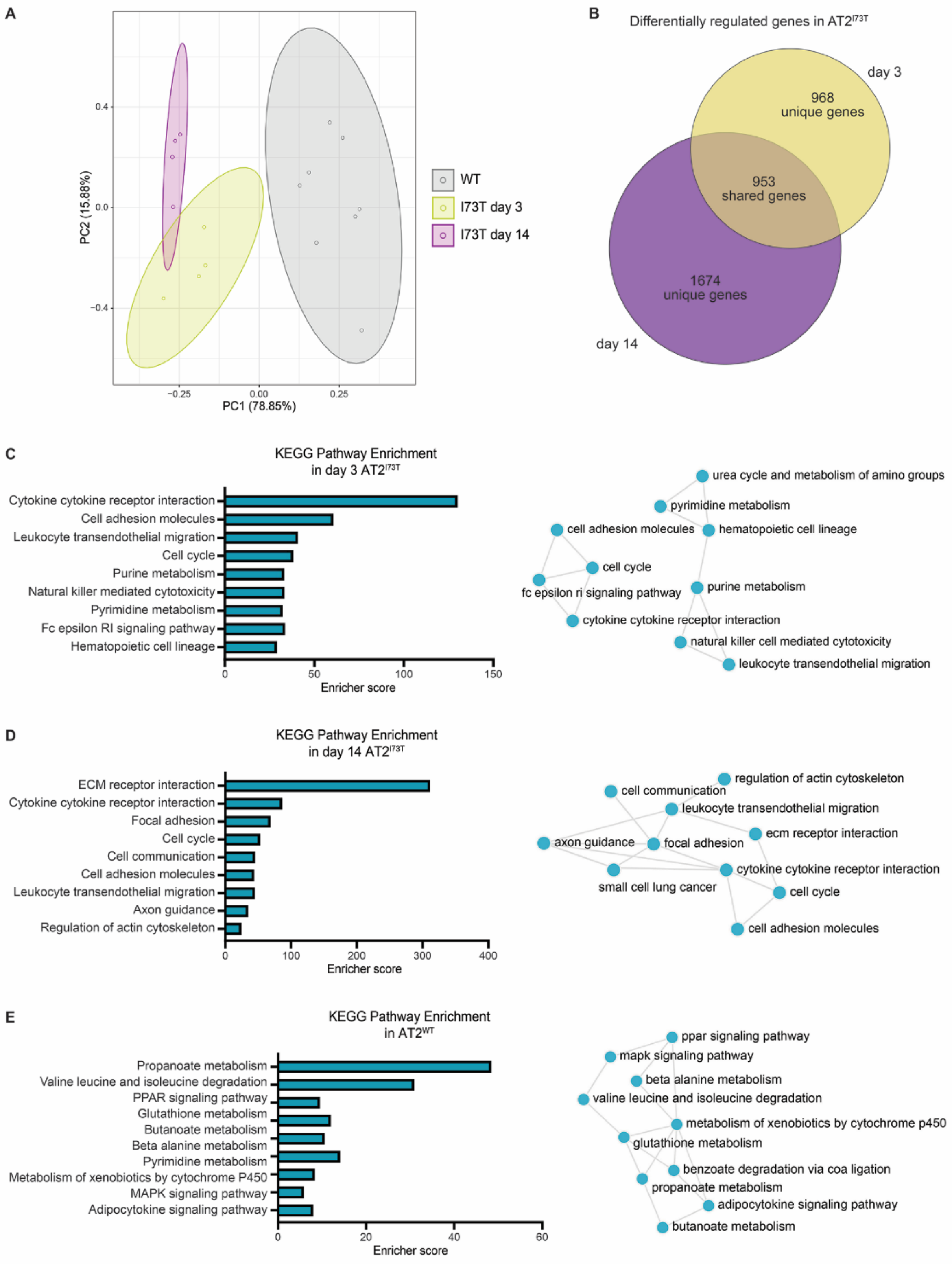
A
PC1 (78.85%)
PC2 (15.88%)
0.4
0.0
−0.4
−0.25
0.00
0.25
WT
I73T day 3
I73T day 14
B
Differentially regulated genes in AT2$^{I73T}$
day 3
968
unique genes
953
shared genes
1674
unique genes
day 14
C
KEGG Pathway Enrichment
in day 3 AT2$^{I73T}$
Cytokine cytokine receptor interaction
Cell adhesion molecules
Leukocyte transendothelial migration
Cell cycle
Purine metabolism
Natural killer mediated cytotoxicity
Pyrimidine metabolism
Fc epsilon RI signaling pathway
Hematopoietic cell lineage
0
50
100
150
Enricher score
urea cycle and metabolism of amino groups
pyrimidine metabolism
cell adhesion molecules
hematopoietic cell lineage
cell cycle
fc epsilon ri signaling pathway
purine metabolism
cytokine cytokine receptor interaction
natural killer cell mediated cytotoxicity
leukocyte transendothelial migration
D
KEGG Pathway Enrichment
in day 14 AT2$^{I73T}$
ECM receptor interaction
Cytokine cytokine receptor interaction
Focal adhesion
Cell cycle
Cell communication
Cell adhesion molecules
Leukocyte transendothelial migration
Axon guidance
Regulation of actin cytoskeleton
0
100
200
300
400
Enricher score
regulation of actin cytoskeleton
cell communication
leukocyte transendothelial migration
ecm receptor interaction
axon guidance
focal adhesion
cytokine cytokine receptor interaction
small cell lung cancer
cell cycle
cell adhesion molecules
E
KEGG Pathway Enrichment
in AT2$^{WT}$
Propanoate metabolism
Valine leucine and isoleucine degradation
PPAR signaling pathway
Glutathione metabolism
Butanoate metabolism
Beta alanine metabolism
Pyrimidine metabolism
Metabolism of xenobiotics by cytochrome P450
MAPK signaling pathway
Adipocytokine signaling pathway
0
20
40
60
Enricher score
ppar signaling pathway
mapk signaling pathway
beta alanine metabolism
valine leucine and isoleucine degradation
metabolism of xenobiotics by cytochrome p450
glutathione metabolism
benzoate degradation via coa ligation
propanoate metabolism
adipocytokine signaling pathway
butanoate metabolism

Figure S1 AT2$^{I73T}$ RNA Seq Analysis: A) Principal component analysis of RNA-seq data derived from AT$^{WT}$ and AT2$^{I73T}$ cells 3- and 14-days post tamoxifen induction. B) Venn diagram highlighting differentially expressed genes when comparing AT2$^{I73T}$ at either 3- or 14-days post induction to AT2$^{WT}$. C) KEGG Pathway enrichment of differentially expressed genes in AT2$^{I73T}$ as compared to the AT2$^{WT}$ 3 days post tamoxifen induction. D) KEGG Pathway enrichment of differentially expressed genes in AT2$^{I73T}$ as compared to the AT2$^{WT}$ 3 days post tamoxifen induction. E) KEGG Pathway enrichment of differentially expressed genes in AT2$^{WT}$ as compared to the AT2$^{I73T}$ at 3- and 14-days post tamoxifen induction. C-E) Analysis was performed using Enricher and presented as both bar graph and network tree.

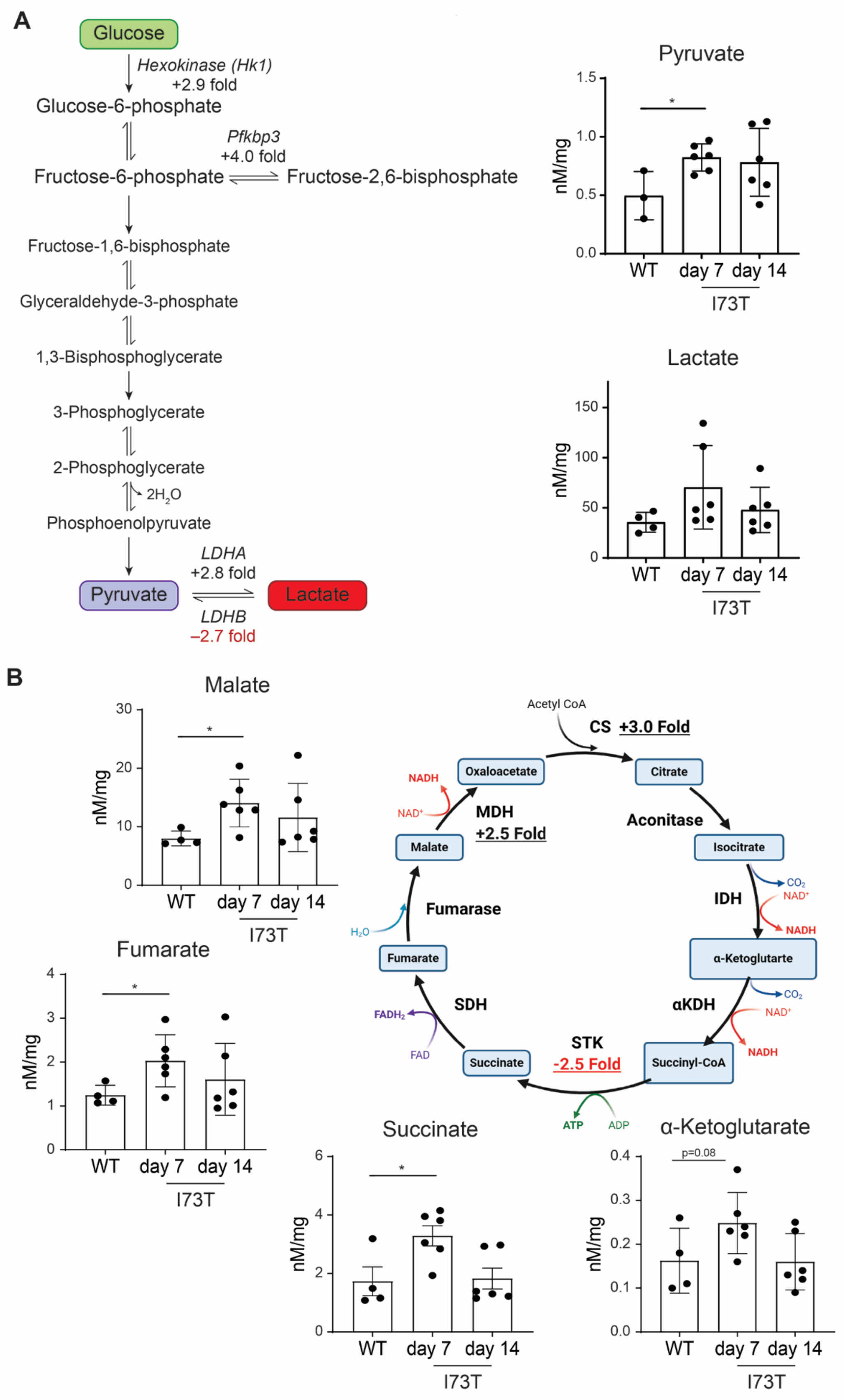

A
Glucose
Hexokinase (Hk1)
+2.9 fold
Glucose-6-phosphate
Pfkbp3
+4.0 fold
Fructose-6-phosphate
Fructose-2,6-bisphosphate
Fructose-1,6-bisphosphate
Glyceraldehyde-3-phosphate
1,3-Bisphosphoglycerate
3-Phosphoglycerate
2-Phosphoglycerate
2H2O
Phosphoenolpyruvate
LDHA
+2.8 fold
Pyruvate
Lactate
LDHB
–2.7 fold
Pyruvate
nM/mg
WT
day 7
day 14
I73T
Lactate
nM/mg
WT
day 7
day 14
I73T
B
Malate
nM/mg
WT
day 7
day 14
I73T
Fumarate
nM/mg
WT
day 7
day 14
I73T
Acetyl CoA
CS +3.0 Fold
Oxaloacetate
Citrate
NADH
NAD+
MDH
+2.5 Fold
Aconitase
Malate
Isocitrate
Fumarase
IDH
CO2
NAD+
NADH
H2O
Fumarate
α-Ketoglutarte
SDH
αKDH
CO2
NAD+
FADH2
STK
FAD
Succinate
-2.5 Fold
Succinyl-CoA
NADH
ATP
ADP
Succinate
nM/mg
WT
day 7
day 14
I73T
α-Ketoglutarate
p=0.08
nM/mg
WT
day 7
day 14
I73T

Figure S2 Organic Acid Quantification in AT2$^{I73T}$: A) Diagram of Glycolysis through the conversion of pyruvate to lactate. Key rate-limiting enzymes and their associated RNA fold changes (AT2$^{I73T}$ at 14-days post tamoxifen induction vs AT2$^{WT}$) are labeled. Bar graphs present the AT2 intracellular concentration of lactate and pyruvate as quantified by mass spectrometry. Ordinary one-way ANOVA testing was performed with statistical significance denoted by $*p < 0.05$. B) Diagram of Krebs Cycle intermediates and associated enzymes. Key enzymes and their associated RNA fold changes are labeled (AT2$^{I73T}$ at 14-days post tamoxifen induction vs AT2$^{WT}$). Bar graphs present the AT2 intracellular concentration of malate, fumarate, succinate, and alpha-ketoglutarate. Ordinary one-way ANOVA testing was performed with statistical significance denoted by $*p < 0.05$.

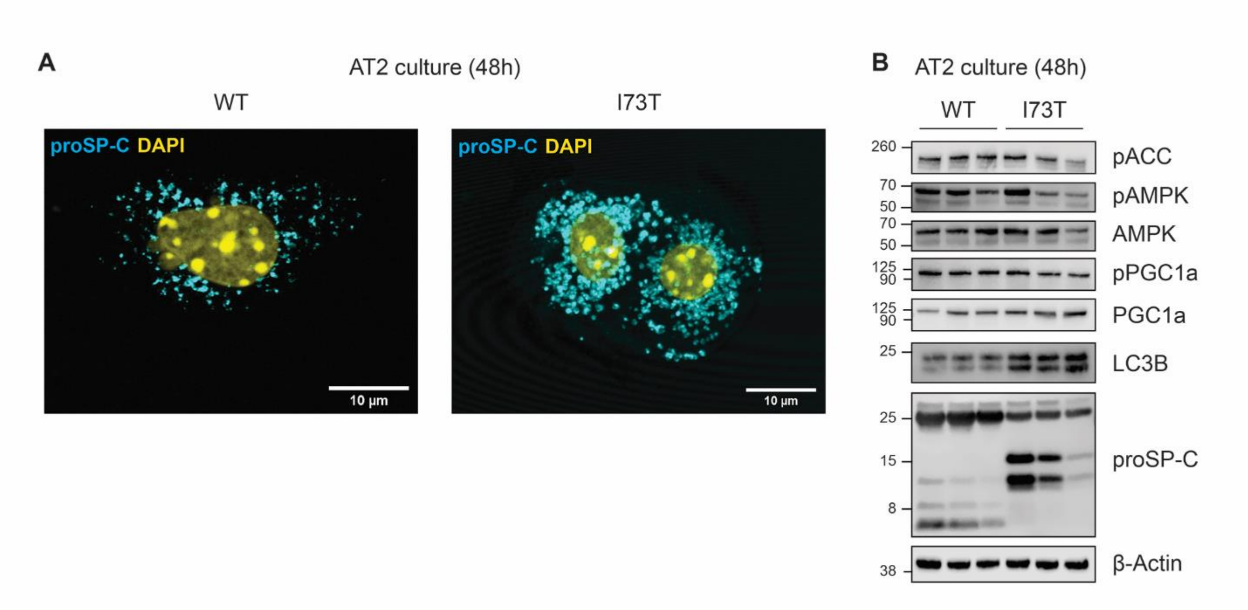


Figure S3 Forty-Eight Hour Culture of Primary Murine AT2s: A) Immunofluorescence staining of $AT2^{WT}$ and $AT2^{I73T}$ 14 days post tamoxifen induction and cultured in 5% DMEM for 48 hours, highlights sustained expression of proSP-C in both mutant and WT AT2s and continued accumulation of I73T isoform. Scale bar 10 μm. B) Western blot analysis of whole cell lysates from 48-hour AT2 culture confirms autophagy dysfunction as seen through accumulation of LC3B as well as the continued accumulation of processing intermediates in $AT2^{I73T}$.

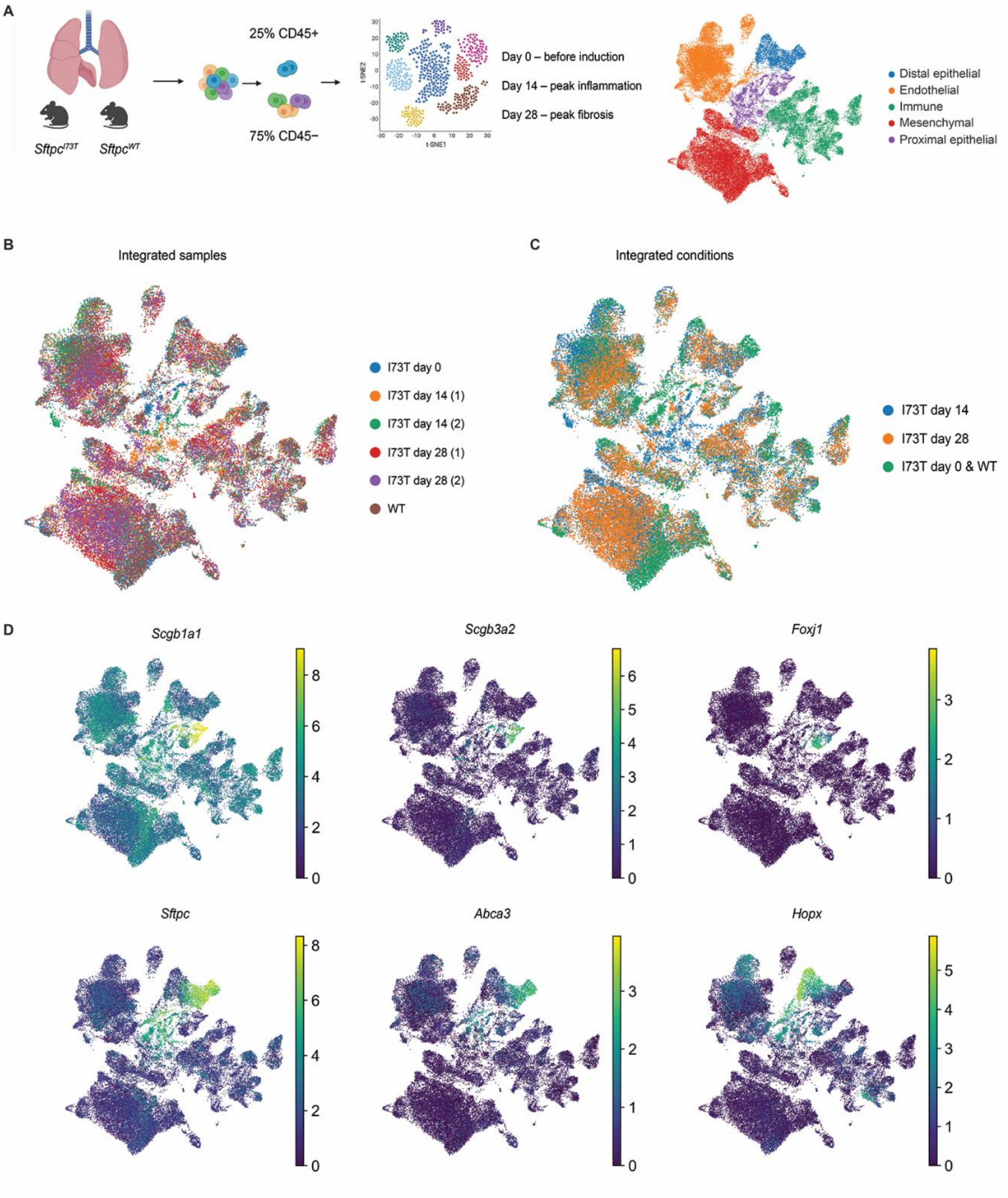
A
25% CD45+
75% CD45−
Sftpc^I73T
Sftpc^WT
Day 0 – before induction
Day 14 – peak inflammation
Day 28 – peak fibrosis
Distal epithelial
Endothelial
Immune
Mesenchymal
Proximal epithelial
B
Integrated samples
I73T day 0
I73T day 14 (1)
I73T day 14 (2)
I73T day 28 (1)
I73T day 28 (2)
WT
C
Integrated conditions
I73T day 14
I73T day 28
I73T day 0 & WT
D
Scgb1a1
Scgb3a2
Foxj1
Sftpc
Abca3
Hopx

Figure S4 Single Cell Analysis of Epithelial Subset in *Sftpc*$^{I73T}$ mice: A) Schematic representation of single cell strategy used to generate GSE234604 consisting of depletion and "add-back spiking of CD45 cells to achieve a final ratio of 25% immune cells and 75% non-immune cells. UMAP projection of the integrated dataset highlighting five major compartments: distal epithelial, proximal epithelial, endothelial, immune, and mesenchyme. B) UMAP projection of the integrated dataset color coded by individual samples C) UMAP projection of the integrated dataset color coded by the model timepoint. Day 0 *SftpcI*$^{73T}$ sample and *Sftpc*$^{WT}$ sample cluster together and are marked by the same color. D) UMAP projection of the integrated dataset highlighting expression of genes used to identify distal and proximal epithelial clusters.

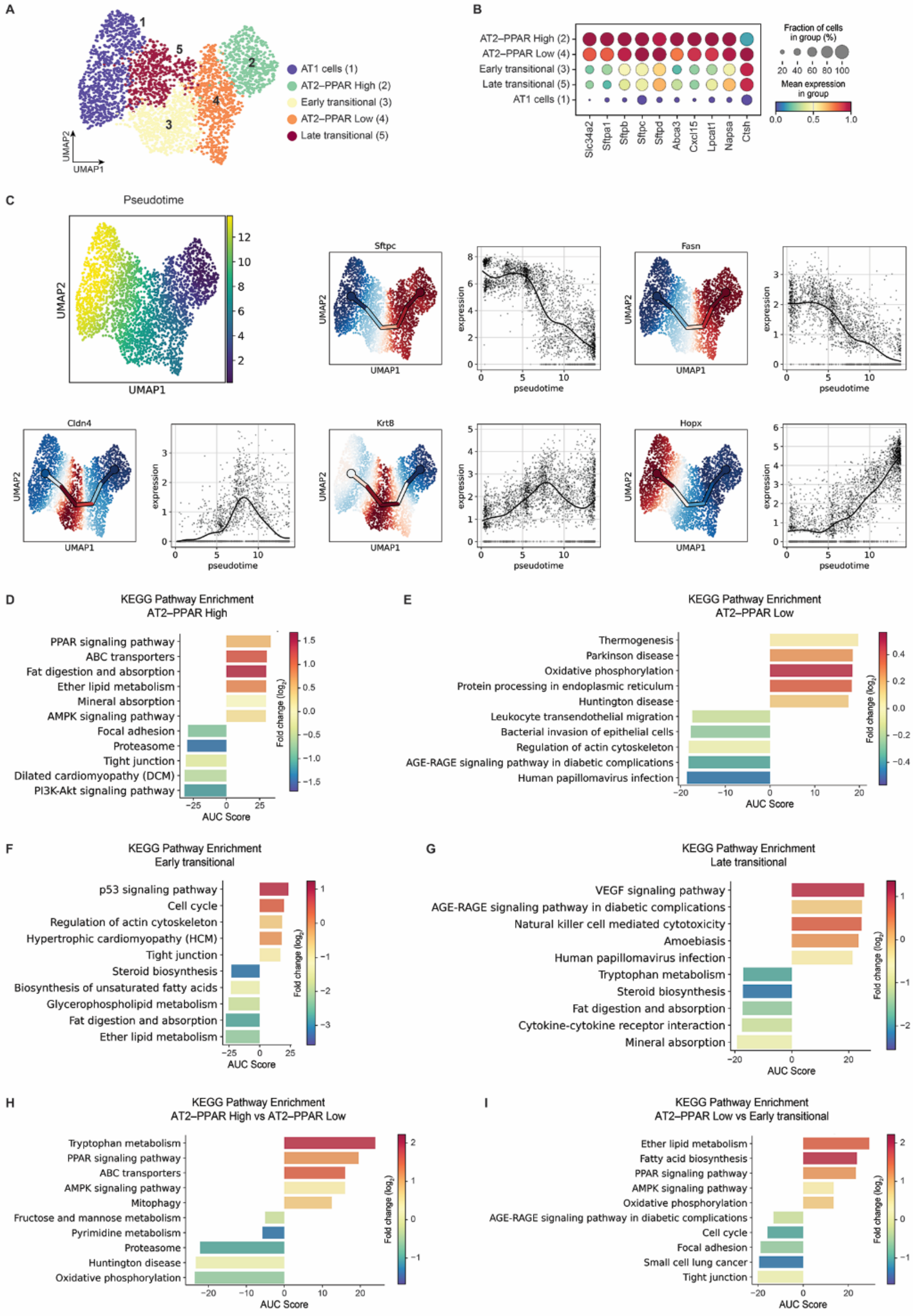
A
AT1 cells (1)
AT2–PPAR High (2)
Early transitional (3)
AT2–PPAR Low (4)
Late transitional (5)
UMAP1
UMAP2
B
AT2–PPAR High (2)
AT2–PPAR Low (4)
Early transitional (3)
Late transitional (5)
AT1 cells (1)
Slc34a2
Sftpa1
Sftpb
Sftpc
Sftpd
Abca3
Cxcl15
Lpcat1
Napsa
Ctsh
Fraction of cells in group (%)
Mean expression in group
C
Pseudotime
Sftpc
Fasn
Cldn4
Krt8
Hopx
expression
pseudotime
D
KEGG Pathway Enrichment
AT2–PPAR High
PPAR signaling pathway
ABC transporters
Fat digestion and absorption
Ether lipid metabolism
Mineral absorption
AMPK signaling pathway
Focal adhesion
Proteasome
Tight junction
Dilated cardiomyopathy (DCM)
PI3K-Akt signaling pathway
AUC Score
Fold change (log2)
E
KEGG Pathway Enrichment
AT2–PPAR Low
Thermogenesis
Parkinson disease
Oxidative phosphorylation
Protein processing in endoplasmic reticulum
Huntington disease
Leukocyte transendothelial migration
Bacterial invasion of epithelial cells
Regulation of actin cytoskeleton
AGE-RAGE signaling pathway in diabetic complications
Human papillomavirus infection
AUC Score
Fold change (log2)
F
KEGG Pathway Enrichment
Early transitional
p53 signaling pathway
Cell cycle
Regulation of actin cytoskeleton
Hypertrophic cardiomyopathy (HCM)
Tight junction
Steroid biosynthesis
Biosynthesis of unsaturated fatty acids
Glycerophospholipid metabolism
Fat digestion and absorption
Ether lipid metabolism
AUC Score
Fold change (log2)
G
KEGG Pathway Enrichment
Late transitional
VEGF signaling pathway
AGE-RAGE signaling pathway in diabetic complications
Natural killer cell mediated cytotoxicity
Amoebiasis
Human papillomavirus infection
Tryptophan metabolism
Steroid biosynthesis
Fat digestion and absorption
Cytokine-cytokine receptor interaction
Mineral absorption
AUC Score
Fold change (log2)
H
KEGG Pathway Enrichment
AT2–PPAR High vs AT2–PPAR Low
Tryptophan metabolism
PPAR signaling pathway
ABC transporters
AMPK signaling pathway
Mitophagy
Fructose and mannose metabolism
Pyrimidine metabolism
Proteasome
Huntington disease
Oxidative phosphorylation
AUC Score
Fold change (log2)
I
KEGG Pathway Enrichment
AT2–PPAR Low vs Early transitional
Ether lipid metabolism
Fatty acid biosynthesis
PPAR signaling pathway
AMPK signaling pathway
Oxidative phosphorylation
AGE-RAGE signaling pathway in diabetic complications
Cell cycle
Focal adhesion
Small cell lung cancer
Tight junction
AUC Score
Fold change (log2)

Figure S5 Pathway Enrichment Analysis of *Sftpc*$^{I73T}$ Distal Alveolar Single Cell Analysis: A) UMAP projection of the integrated distal alveolar dataset identifying five cluster. B) Dot plot of AT2 associated gene expression in the five clusters identified in the distal alveolar UMAP C) Pseudotime analysis of the distal alveolar UMAP and individual gene pseudotime plots that represent some key markers of the AT2-to-AT1 transition D) KEGG Pathway enrichment analysis identifying the top 5 pathways that are significantly enriched and decreased in the AT2-PPAR high cluster as compared to the other 4 clusters. E) KEGG Pathway enrichment analysis identifying the top 5 pathways that are significantly enriched and decreased in the AT2-PPAR low cluster as compared to the other 4 clusters. F) KEGG Pathway enrichment analysis identifying the top 5 pathways that are significantly enriched and decreased in the Early Transitional cluster as compared to the other 4 clusters. G) KEGG Pathway enrichment analysis identifying the top 5 pathways that are significantly enriched and decreased in the Late Transitional cluster as compared to the other 4 clusters.

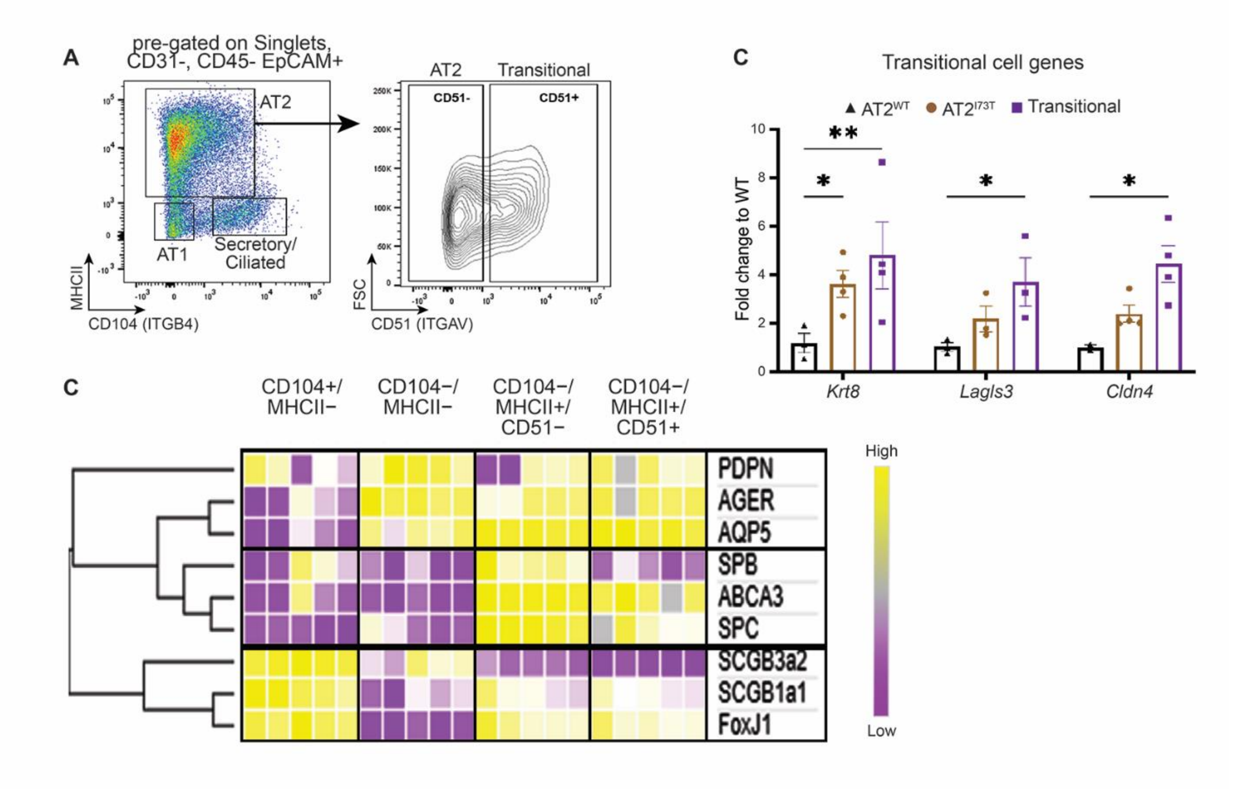


Figure S6 Isolation of transitional AT2 cells: A) Gating of flow cytometry strategy used to isolate transitional AT2 cells, not shown is the pre-gating for live cells that are CD45$^{neg}$, CD31$^{neg}$, and Epcam$^{+}$. B) Heatmap of QPCR gene expression fold changes in distal alveolar identity genes derived from the various population isolated through the transitional AT2 gating strategy. All genes are normalized to the mean expression of the individual gene in the four populations. Hierarchical clustering using Euclidean distance was performed on heatmap rows. C) Bar graph of select transitional cell identity gene expression fold change in CD51- and CD51+ populations derived from WT and I73T AT2s. Ordinary one-way ANOVA testing was performed with statistical significance denoted by *p < 0.05 ** p<0.005.

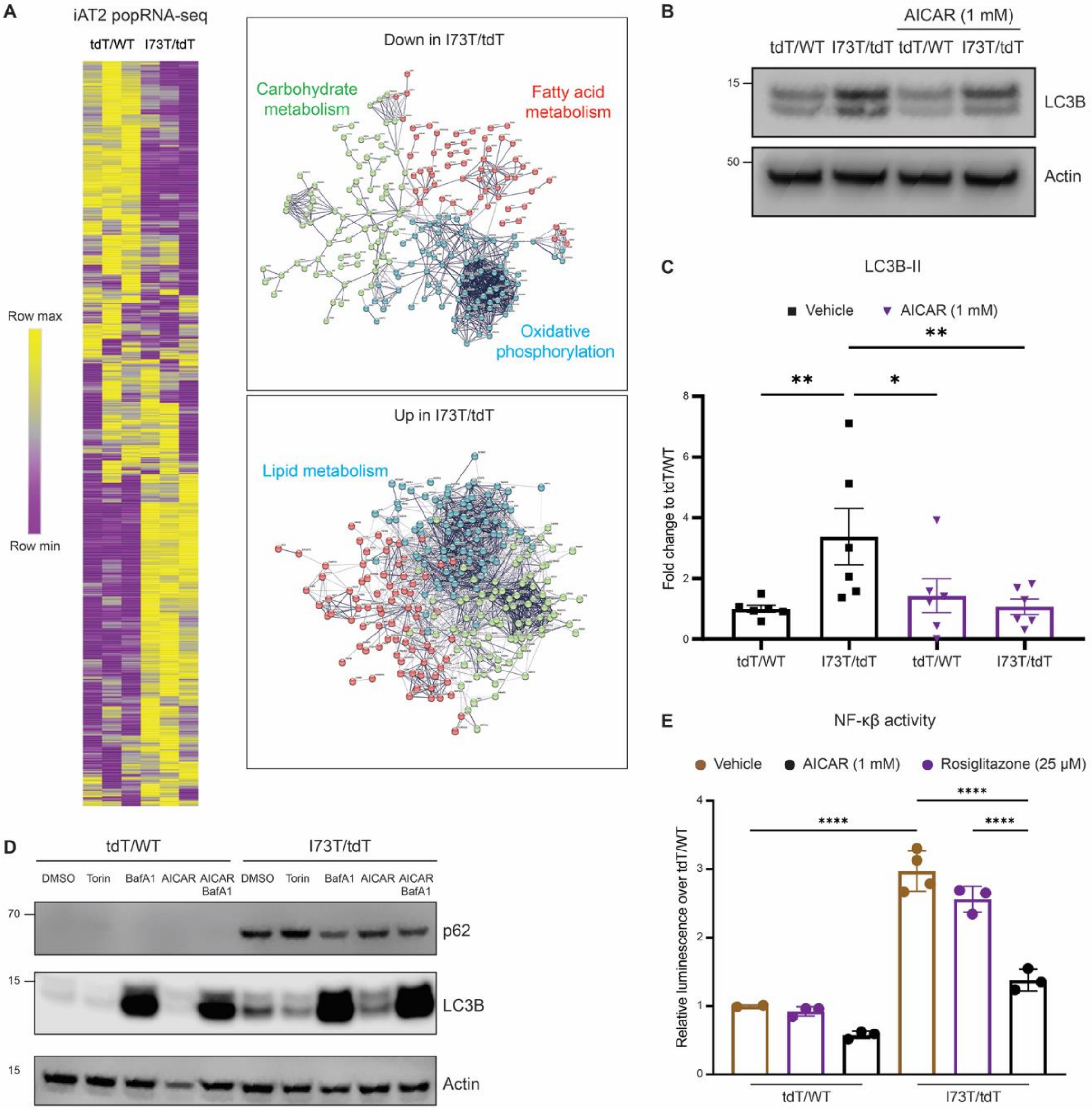

A
iAT2 popRNA-seq
tdT/WT
I73T/tdT
Row max
Row min
Down in I73T/tdT
Carbohydrate metabolism
Fatty acid metabolism
Oxidative phosphorylation
Up in I73T/tdT
Lipid metabolism
B
AICAR (1 mM)
tdT/WT
I73T/tdT
tdT/WT
I73T/tdT
LC3B
Actin
C
LC3B-II
Vehicle
AICAR (1 mM)
Fold change to tdT/WT
tdT/WT
I73T/tdT
tdT/WT
I73T/tdT
D
tdT/WT
I73T/tdT
DMSO
Torin
BafA1
AICAR
AICAR BafA1
p62
LC3B
Actin
E
NF-κβ activity
Vehicle
AICAR (1 mM)
Rosiglitazone (25 μM)
Relative luminescence over tdT/WT
tdT/WT
I73T/tdT

Figure S7 I73T iAT2 Autophagy Defect and Inflammatory Activation is Ameliorated via AMPK Agonism : A) String DB analysis of differentially (significant by FDR) expressed genes associated with metabolic pathways in GSE160801 identifies several altered metabolic pathways including carbohydrate metabolism, fatty acid metabolism, lipid metabolism, and oxidative phosphorylation. B) Representatives immunoblot of LC3B expression in corrected and mutant iAT2s after overnight 1mM AICAR challenge. C)  Densitometry quantification of multiple replicates (n=6) normalized to the mean densitometry of the corresponding corrected mean included in each immunoblot. D) Immunoblot of autophagy markers p62 and LC3B from cell lysates of corrected and mutant iAt2s after overnight challenge with either 5μM Torin, 50nM Bafylomicin A, 1mM AICAR, or combination. E) After lentiviral transduction of mutant and corrected iAT2s with NF-κB-luc-GFP lentiviral construct cells were challenged with 1mM AICAR or 25mM Rosiglitazone overnight and bioluminescence quantification shows increased luciferase activity in mutant cells that is significantly decreased after AMPK agonism. Ordinary one-way ANOVA testing was performed with statistical significance denoted by *$p < 0.05$ ** $p<0.005$.
